# Rational design of protein-specific folding modifiers

**DOI:** 10.1101/2020.04.28.064113

**Authors:** Anirban Das, Anju Yadav, Mona Gupta, R Purushotham, Vishram L. Terse, Vicky Vishvakarma, Sameer Singh, Tathagata Nandi, Kalyaneswar Mandal, Shachi Gosavi, Ranabir Das, Sri Rama Koti Ainavarapu, Sudipta Maiti

## Abstract

Protein folding can go wrong *in vivo* and *in vitro*, with significant consequences for the living cell and the pharmaceutical industry, respectively. Here we propose a general design principle for constructing small peptide-based protein-specific folding modifiers. We construct a ‘xenonucleus’, which is a pre-folded peptide that resembles the folding nucleus of a protein, and demonstrate its activity on the folding of ubiquitin. Using stopped-flow kinetics, NMR spectroscopy, Förster Resonance Energy transfer, single-molecule force measurements, and molecular dynamics simulations, we show that the ubiquitin xenonucleus can act as an effective decoy for the native folding nucleus. It can make the refolding faster by 33 ± 5% at 3 M GdnHCl. In principle, our approach provides a general method for constructing specific, genetically encodable, folding modifiers for any protein which has a well-defined contiguous folding nucleus.

## Introduction

Protein sequences have evolved to fold efficiently^1–6^. Still, folding can go wrong *in vivo*^7^, initiating a large number of diseases^8–12^. It can also do so *in vitro*^13,14^, with significant consequences for the food and pharmaceutical industries^15–18^. The cell has evolved elaborate chaperone machinery to steer folding^19–23^, but designing artificial protein-specific folding modifiers has remained a challenge^24–27^. Here we demonstrate a general design principle for constructing small peptide or peptidomimetic protein-specific folding modifiers. It is based on making a ‘xenonucleus’, which is a separate pre-folded peptide (or artificial peptide-mimic) that resembles the ‘folding nucleus’ of a protein.

Rates of protein folding are evolutionarily dictated by its sequence^28–30^. Folding occurs on a multidimensional pathway, but many small proteins fold through a reasonable set of well-defined conformations, often termed ‘foldons’^31–34^. The lowest free energy path through which an unfolded protein travels, moving through this set of conformations to reach the folded state, is known as the folding pathway. For many proteins, one region folds first, and its interaction with other regions helps the whole protein fold. This region is termed as the ‘folding nucleus’^35–40^. It is believed that the folding nucleus can transiently adopt a native-like conformation, even in the unfolded form of the protein^30,41–44^. The nucleus itself may fold and unfold many times before the whole protein folds. Nucleus formation is frequently, though not always^45,46^, the step that determines the folding kinetics. This suggests that holding the nucleus stable in the native-like conformation should speed up protein folding. Indeed, folding has been shown to speed up in experiments where the nucleus has been held stable through chemical means^47^.

Here we take this logic further, and ask whether an unfolded protein will be able to recognize and fold using a pre-folded but separate ‘xenonucleus’ which can act as a decoy. A short stretch of a peptide with a sequence identical to that of the folding nucleus (or the ‘minimalistic mimic’) may not fold by itself. However, introducing appropriate structural constraints between the distal residues, such as a disulphide links, can hold it in a near folded conformation. The concentration of such peptides in the solution must be sufficient so that a protein will make multiple encounters with such objects before it has the time to fold using its own nucleus. Whether a protein that folds in such a manner will remain folded with the xenonucleus in place and with its own nucleus orphaned, cannot be predicted a priori. In any case, this will provide an alternative pathway for folding, and may be expected to affect the folding kinetics. In that sense, the xenonucleus will act as a micro-chaperone, sculpting a new folding pathway for the protein.

We test this strategy on ubiquitin, which is a small, 76-residue protein that plays a primary role in protein recycling by living cells^48^. Ubiquitin is an appropriate model, since its folding pathway has been determined by experiments^49–55^, and also by full atomic scale MD simulations^56^ which agree with those experiments. We construct a ‘xenonucleus’ for ubiquitin (residues 1 to 17, which is the segment that is known to fold first), and conformationally constrain it by introducing a pair of disulphide linked cysteines at the two termini. We then study the kinetics and the thermodynamics of the folding of ubiquitin in the presence of this xenonucleus, using several different biophysical tools. The primary questions we ask are whether the kinetics and the thermodynamics of folding are affected by the xenonucleus (and other control compounds), and if so, whether the interaction with the xenonucleus happens at the site where the native nucleus is usually nestled. The results of our stop-flow kinetics, time resolved Forster Resonance Energy transfer, NMR, and coarse grain simulation studies show that kinetics of ubiquitin folding can indeed be made faster by a suitable xenonucleus interacting at the appropriate site.

## Results

### Refolding kinetics

Ubiquitin has been widely studied for its thermodynamic and mechanical stability. Previous mutational studies on Ubiquitin, especially using the F45W mutant, has established tryptophan fluorescence as a valuable probe for studying its folding. Here, we measured the refolding kinetics of Ubiquitin (F45W) in presence and in absence of a 19 residue nucleus mimic. It has the amino acid sequence **C**-MQIFVKTLTGKTITLEV-**C**, which is the same as residues 1 to 17 of Ubiquitin, except for the terminal cysteines. It has a disulphide bridge between the termini, and will henceforth be called the ‘stapled xenonucleus’. We use a three syringe stopped-flow fluorescence instrument (SFM300, Biologic, see Methods) to change the GdnHCl concentration from 4.25 M to a series of lower concentrations in less than 6 ms. We measure the change in fluorescence intensity of the Trp residue as a function of time as a reporter for the progress of folding (or unfolding). The unfolded state of ubiquitin F45W has a higher quantum yield and has an emission maximum at 360 nm, but when it goes to the folded form, the emission blue-shifts to 340 nm. The quantum yield of fluorescence also goes down (it is speculated that a backbone carboxyl oxygen may be quenching the fluorescence)^57,58^. Hence, as folding proceeds, the overall fluorescence signal at 360 nm decreases. We have plotted the change in the fluorescence intensity as a function of time for ubiquitin F45W, with (Figure 1 (A), red) and without (Figure 1 (A), blue) the xenonucleus. The data are fitted with a two-component exponential decay function:

**Figure 1.**
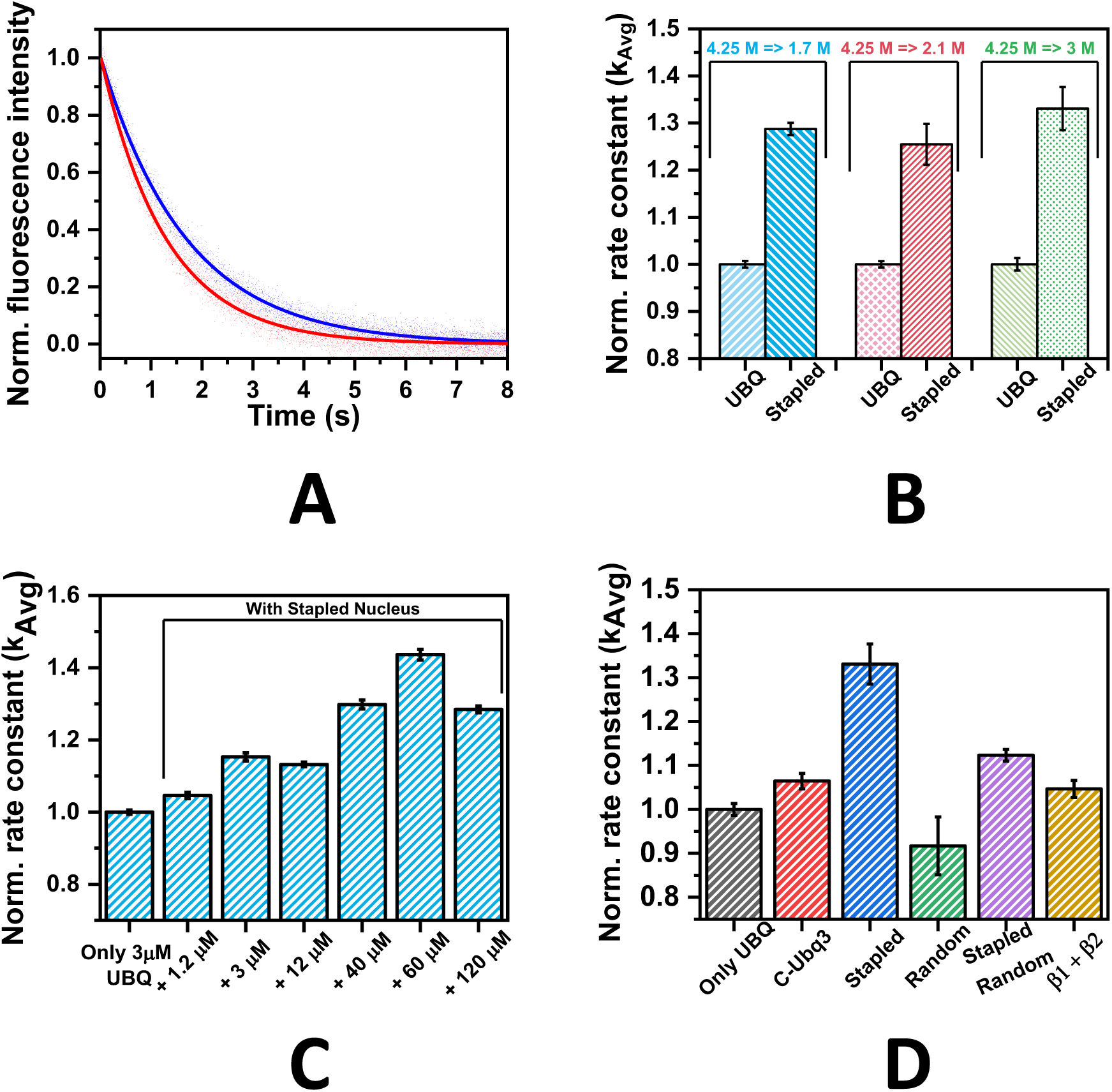
(A) Folding of Ubiquitin (Ubq F45W) as reported by Trp fluorescence in a stop-flow kinetics experiment. Ubiquitin alone (black) and in presence of the stapled xenonucleus (red), at a final GdnHCl concentration of 3 M GdnHCl starting from a GdnHCl concentration of 4.25 M. (B) folding rate constants at final GdnHCl concentrations of 1.7 M (blue), 2.1 M (red) and 3 M (green). (C) Xenonucleus concentration dependence of the folding rate constant (final [GdnHCl] = 1.7 M) (D) folding rate constants in presence of different peptides (final [GdnHCl] = 3 M).

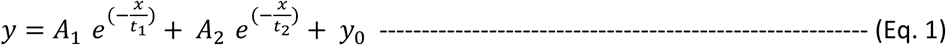

Here, A_i_ and t_i_ and y_0_ are the amplitude, the average lifetime of the i-th component, and the initial signal at time t = 0, respectively. The corresponding rate constants are calculated subsequently using the relation, k_i_ = 1/t_i_ (i = 1, 2) and the average rate constant is the amplitude weighted average of the two rate constants, k_Avg_ = {(A_1_*k_1_ + A_2_*k_2_)/(A_1_ + A_2_)}. Clearly, in presence of the xenonucleus for the 3 M GdnHCl end-point (which is the mid-point for the GdnHCl mediated unfolding), the refolding rate (average k_folding_) becomes 33 ± 5% faster than that observed for ubiquitin alone. In addition, this effect is consistently found at all the different end-point GdnHCl concentrations, namely (i) 1.7 M (1.29 ± 0.01 times), and (ii) 2.1 M (1.25 ± 0.04 times), as shown in Figure 1 (B).

### Refolding kinetics as a function of the xenonucleus concentration

We also perform the refolding experiment as a function of the concentration of the xenonucleus peptide, for a fixed dilution of the GdnHCl concentration from 4.25 M to 1.7 M. The concentration of the protein is kept fixed at 3 µM as the peptide concentration is increased from 0 µM to 120 µM. Figure 1 (C) shows the extracted values of the average k_folding_, normalized w.r.t. the kinetics observed at zero peptide concentration. It shows that the rate of refolding increases with the concentration of the peptide, though this effect appears to saturate above 60 µM.

### Sequence dependence of the effect of the xenonucleus

We probe the specificity of the interaction between the xenonucleus peptide and the protein by repeating the refolding experiment (for GdnHCl dilution from 4.25 M to 3 M) with many different variants of the stapled xenonucleus. The peptides used were Cys-Ubq3 (C-MQIFVKTLTGKTITLEV), Random peptide (same peptide, randomized sequence: TGVT**C**MVFIELLQKTTKI), Stapled Random peptide (**C**-TGVTMVFIELLQKTTKI-**C**, with a disulphide bridge between the termini), and cleaved Cys-Ubq3 fragments (Cys-Ubq1: **C**-MQIFVKTL and Ac-Ubq2: **Ac**-GKTITLEV). In all the cases, we see some increase in the refolding rate (Figure 1 (D)), but the increase for all the other peptides is substantially less than that with the stapled xenonucleus.

### Effect of the xenonucleus on the thermodynamic stability of the protein

The faster kinetics of folding implies that the xenonucleus stabilizes the folded state with respect to the unfolded state. Alternatively, it may effectively change only the free energy barrier between the two states, acting as a ‘folding catalyst’. We test this by performing a steady state experiment measuring the fluorescence intensity at 360 nm as a function of increasing GdnHCl concentration. Figure 2 (A) and Figure S12 (A) show that the fluorescence intensity, as expected, goes up as the protein (at 10 µM) is unfolded by the addition of GdnHCl. The data are fitted with the sigmoidal function related to the chemical denaturation of a protein:

**Figure 2.**
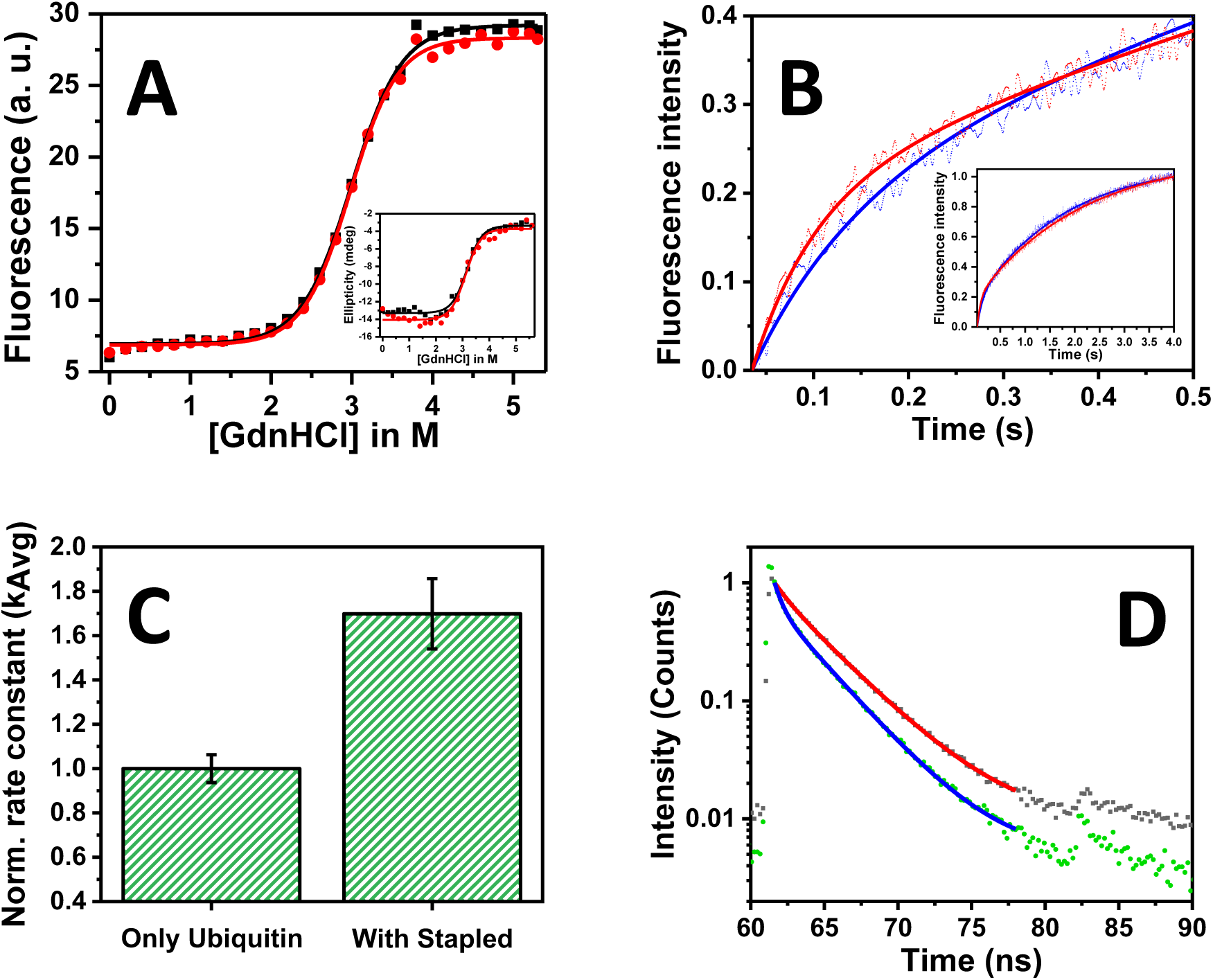
(A) Steady state unfolding of Ubiquitin by GdnHCl in the absence (black) and in the presence (red) of the stapled xenonucleus, as measured by Trp fluorescence at 360 nm, and (inset) by circular dichroism at 222 nm. (B) Unfolding kinetics and (C) normalized unfolding rate constants of Ubiquitin in absence (blue) and presence (red) of the stapled xenonucleus (final [GdnHCl] = 3 M) (D) Time resolved fluorescence decay of Rh110-labelled Ubiquitin in presence of unlabelled xenonucleus. Data (black) and fit (red). Same in presence of Cy5-labelled xenonucleus, data (green) and fit (blue).

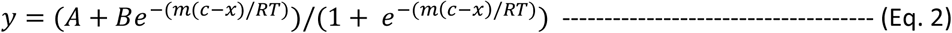

Here x = concentration of denaturant, c = [Denaturant]_50%_ i.e. the concentration of the denaturant at the midpoint of the unfolding transition, m is the slope of the transition, i. e.

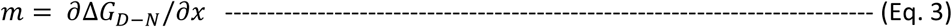

and A and B are constants used as fitting parameters. In a separate experiment, the protein (10 µM) is similarly titrated against GdnHCl but in the presence of the xenonucleus (40 µM) (Figure 2 (A) and Figure S12 (B)). It is obvious from Figure 2 (A) that neither the slope m nor the C_m_ value changes significantly in the presence of the xenonucleus.

We also separately probe this process by measuring the circular dichroism (CD) signal at 222 nm as a function of increasing GdnHCl concentration. A change in the tryptophan fluorescence gives information about the change in the local environment of the tryptophan, while the circular dichroism signal provides information about the global change in the secondary structure. CD results are found to be similar to those obtained from the fluorescence experiment (Figure 2 (A), inset; Figure S13 (A) and (B)).

### Kinetics of ubiquitin unfolding

An increase in the folding kinetics without a change in the equilibrium would imply that the unfolding kinetics also becomes faster. This can be probed by a kinetic unfolding experiment. Ubiquitin (30 µM) is equilibrated at 1 M GdnHCl, and then the protein is unfolded by rapidly mixing it with 4.8 M GdnHCl in 20 mM pH 7.5 PBS (the final GdnHCl concentration is 3 M). In this experiment, the buffer in the second syringe is replaced with the stapled xenonucleus (at 200 µM concentration in pH 7.5 PBS). After the mixing, the final concentrations of the protein and the peptide become 3 μM and 60 μM, respectively. Change in the fluorescence intensity is recorded as a function of time and the data is fitted with a two-exponential function, Eq. 1. Figure 2 (B) and (C) show that the kinetics of unfolding indeed becomes faster in the presence of the xenonucleus, and this increase in k_unfolding_ is somewhat more than the increase of k_folding_ in 3 M GdnHCl.

### Direct interaction of the protein with the xenonucleus

While both the refolding and the unfolding kinetics show an effect of the xenonucleus, it does not establish that the effect is due to a direct interaction of the xenonucleus with the protein. To address this question, we use the Förster Resonance Energy Transfer (FRET) technique. We covalently label the protein (Ubiquitin F45W) with Rhodamine110-NHS ester (Rh110-Succinimidyl ester, the donor) using any available primary amine, and the xenonucleus with Cyanine5-NHS ester (Cy5-Succinimidyl ester, the acceptor) at its N-terminal. The first labelling was performed at a ratio of 1:10 (dye:protein), so that the probability of multiple labelling of the same protein would be low. A decrease in the donor fluorescence lifetime in the presence of the acceptor labelled peptide would establish a molecular interaction between the protein and the xenonucleus. We measure the fluorescence lifetime of the folded protein at 3 M GdnHCl with and without the external peptide, using a time-correlated single-photon counting (TCSPC) set-up as described in the methods section. Figure 2 (D) shows the lifetime decay traces of Rh110-labelled Ubiquitin F45W (3 μM) equilibrated with the unlabelled xenonucleus (120 μM final concentration) and with the Cy5-labelled xenonucleus (3 μM final concentration), at a GdnHCl concentration of 3 M. The lifetime traces are fitted with a 2-component exponential decay function using the Fluofit software (PicoQuant, Germany). The average lifetime for Rh110-Ubiquitin in absence of peptide is 3.56 ± 0.034 ns, in presence of unlabelled xenonucleus is 3.36 ± 0.019 ns, and in presence of labelled xenonucleus is 2.85 ± 0.023 (see SI, Figure S14). This shortening of the lifetime indicates energy transfer and implies that the xenonucleus directly interacts with the protein. The individual components are presented in a table (see SI, Table S5).

### Site of interaction

We then probe whether the site of interaction of the xenonucleus is the same as that of the native nucleus. We employ nuclear magnetic resonance (NMR) spectroscopy to address this question. Each amide backbone resonance of a protein shows characteristic chemical shifts depending on its chemical environment, and the chemical shifts are sensitive to any changes in the environment. We expect the binding interface of ubiquitin (UBQ) to detect the minor changes in the chemical environment when the xenonucleus binds. We have recorded the ^15^N-edited two-dimensional (2D) Heteronuclear Single Quantum Coherence (HSQC) spectra of the ^15^N isotopically labelled UBQ (100 µM) at 1.7 M GdnHCl with and without the unlabelled xenonucleus (Stapled Cys-Ubq3-Cys peptide, 100 µM). We note that higher GdnHCl concentrations produced artefacts In NMR. The effects of xenonucleus on the UBQ are detected by these experiments. Figure 3 (A) shows an overlay of the HSQCs of free UBQ and UBQ/peptide complex at 1.7 M GdnHCl. Figure 3 (B) shows the Chemical Shift Perturbation (CSP), defined as ((dN/5)^2+(dH)^2))^1^/^2^, where dH(dN) is the chemical shift difference of the proton (^15^N) between ubiquitin and ubiquitin/xenonucleus complex. Interestingly, the CSPs are small compared to typical protein/peptide complexes, which indicates that the xenonucleus peptide is binding to UBQ in the β1-β2 conformation and at the same interface, such that the chemical shifts of peptide bound UBQ is close to its native state. Figure 3 (C) shows the CSPs mapped on the structure of UBQ. The significant CSPs are specific to the β1-β2 binding site, confirming that the xenonucleus is binding at the same interface where β1-β2 binds. Maximum CSPs are observed for the residues Val26, Lys29 and Lys33 present in the first helical region (α1) and again for the residues Leu67, Leu69 and Leu71 present in the last β-strand (β5) respectively. Interestingly, these are the residues which are found to be in the closest proximity of the beta1-beta2 in the native form of the protein (Table S4, in the SI). The maximum CSP observed for Val26 also correlates well with its close distance with the beat1-beta2 (Val17 - 4.24 Å between the terminal carbon of both valines and Ile3 - 4.92 Å between the terminal carbons). A few additional CSPs are observed in the β1-β2 region of the protein (e. g. Glu16 - 2.67 Å between terminal Nitrogen of Lys29 and backbone oxygen of Glu16, Table S4), suggesting that the xenonucleus is competing and displacing the β1-β2 region to a non-native environment. And probably because of the same reason, CSP for the Thr22 residue present at the start of the helix (α1) is also found to be quite substantial. We repeat the experiments by mixing the UBQ and the xenonucleus at 0 M GdnHCl, and compare with free UBQ spectra at 0 M GdnHCl concentration (shown in the SI, Figure S16). The CSPs at 0 M GdnHCl are insignificant compared to that observed at 1.7 M GdnHCl, which indicates that the xenonucleus does not interact with the protein in the folded state.

**Figure 3.**
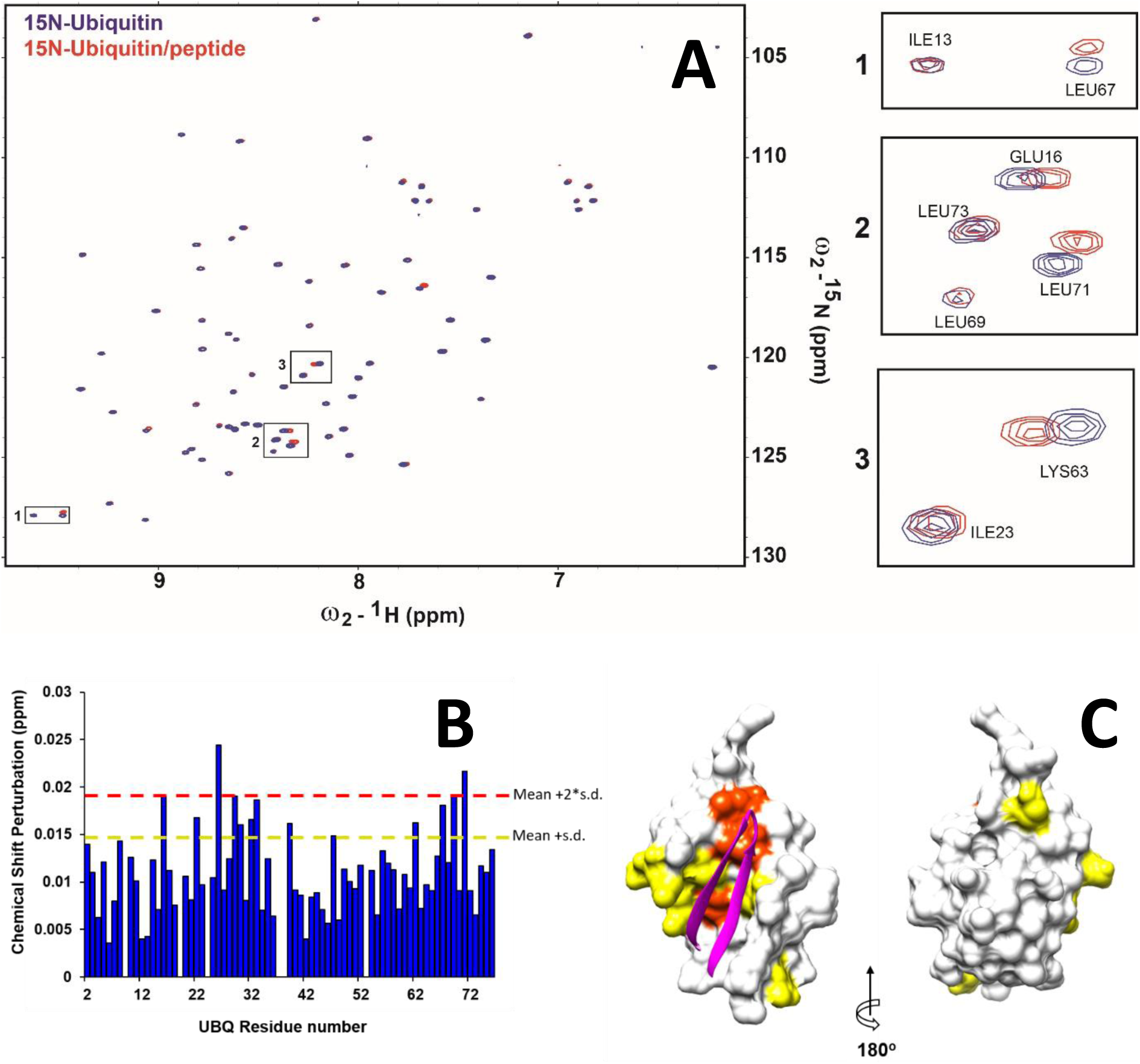
(A) Overlay of HSQCs of free UBQ (blue) and UBQ/xenonucleus complex (red) at 1.7 M GdnHCl, (B) Chemical Shift Perturbation [((dN/5)^2+(dH)^2))^1^/^2^] between UBQ and UBQ/peptide solution at 1.7 M GdnHCl, (C) Chemical Shift Perturbation [((dN/5)^2+(dH)^2))^1^/^2^] mapped to the structure of UBQ. The β1-β2 region is shown in magenta ribbon, while the rest is shown as surface. CSPs > mean + s. d. are coloured in yellow and CSPs > mean + 2 s. d. are coloured in orange.

It is likely that this site-specific interaction is driven both by the overall conformation of the peptide, as well as the local stereospecific interactions of individual residues with their surroundings. Our experiments so far do not distinguish between these two effects. Therefore, we synthesized the D-amino acid analogue of the xenonucleus (‘Stapled Cys-Ubq3-Cys’), namely the ‘stapled D-peptide’ (amino acid sequence: cmqifvktltGktitlevc, with a disulphide bridge between the two termini). This should preserve the secondary structure but would tend to adopt a conformation which is the mirror image of the original xenopeptide. The sidechains of the stapled D-peptide may not have similar conformations as the L-isomorph, so the interaction is expected to be less. We performed ^15^N-edited 2D HSQC experiments of the ^15^N-labelled UBQ (100 µM) at 1.7 M GdnHCl with and without the unlabelled stapled D-peptide (100 µM). Figure S17 shows the CSPs between the UBQ and the UBQ/peptide (Stapled D-peptide) complex at 1.7 M GdnHCl. The chemical shift differences between the ^15^N-edited HSQC of UBQ and the UBQ/peptide complex tend to suggest that the D-peptide also interacts with the same area. However, the amplitude of the CSPs indicates that the affinity of D-peptide is lower than that of the L-peptide, suggesting that the interaction is partly determined by stereochemistry, and partly by the sequence of the xenonucleus.

### Interaction between neighbouring ubiquitins in the presence of the xenonucleus

If the xenonucleus occupies the space that originally was occupied by the β1-β2 segment, then that part should be free to interact with other neighbouring ubiquitin molecules. Although the free beta1-beta2 segment is not structurally constrained, it may act as the nucleus for another ubiquitin molecule. We probe this possibility using Atomic Force spectroscopy (AFM) of a ubiquitin nonamer, where each ubiquitin molecule is covalently connected through a peptide link to the next ubiquitin. The protein is equilibrated at 1.7 M GdnHCl with the xenonucleus for 1 hour, and then it is unfolded by pulling with an AFM tip along the N to C direction (at a pulling speed of 400 nm/s). Control experiments are performed on a solution of ubiquitin (no xenonucleus) equilibrated at the same GdnHCl concentration. Figure 4 shows the unfolding traces for ubiquitin (Figure 4 (A)), and of ubiquitin with the xenonucleus (Figure 4 (B)). The force-versus-extension (F-X) traces are then fitted with the worm-like chain (WLC) model of polymer elasticity. The rupture force and the corresponding contour lengths obtained from the data are plotted in the scattered plot shown in Figure 4 (C). Figure 4 (D) and (E) show the unfolding traces obtained at 0 M GdnHCl. Figure 4 (F) shows the corresponding scatter plot of the rupture force vs contour lengths. These results clearly show that in the presence of 1.7 M GdnHCl, there is a decrease in the rupture force, from 200 pN to about 170 pN and occasional increase in the contour length of a single ubiquitin (∼24 nm). The decrease in the rupture force is expected, as the addition of GdnHCl destabilizes the folded protein. In the presence of the xenonucleus, we observe a further decrease in the unfolding force (∼100 pN), and a much more frequent increase in the contour length. No effect of the xenonucleus is observed when the protein is mixed with the peptide at 0 M GdnHCl.

**Figure 4.**
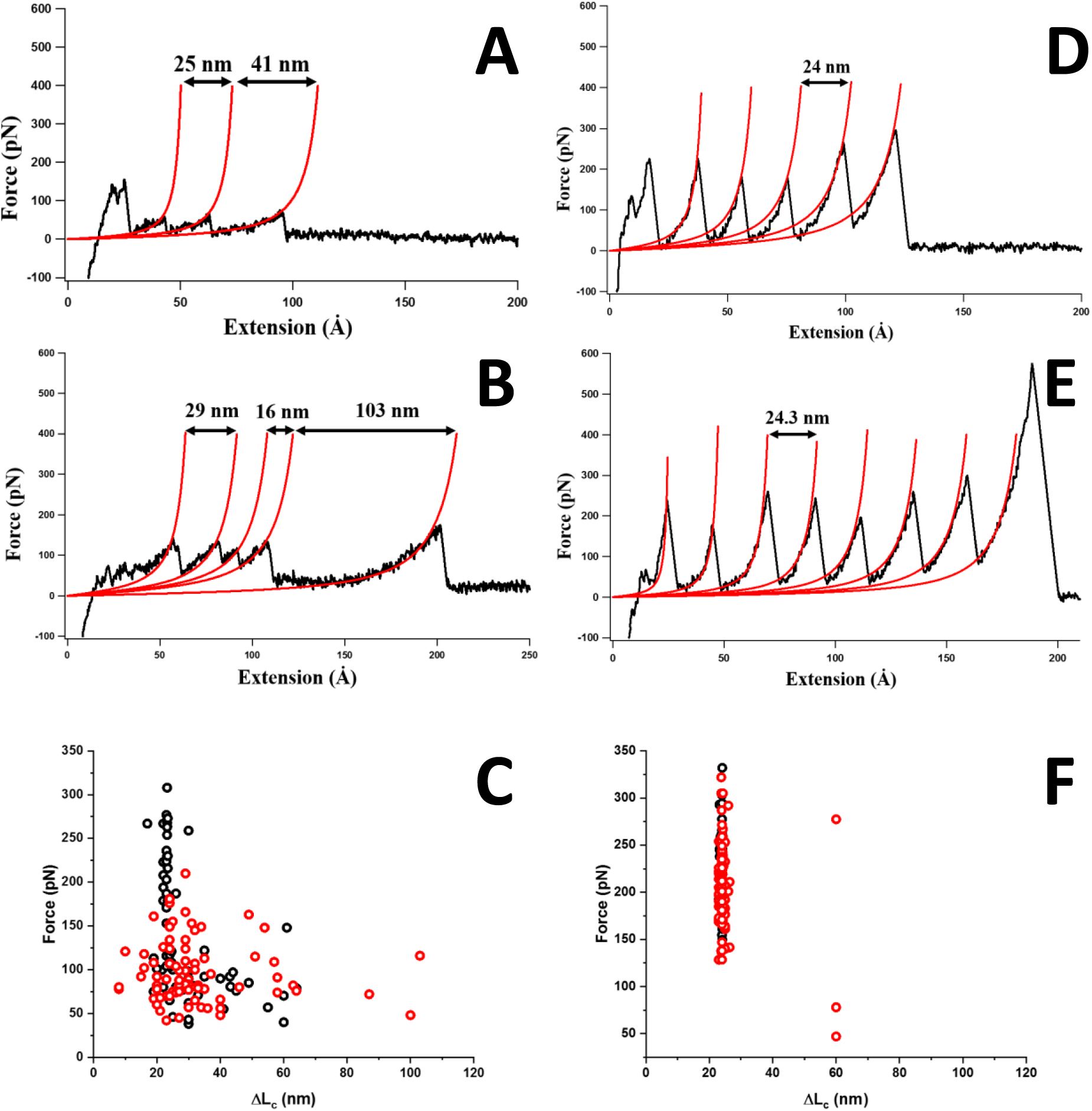
AFM force-pulling traces of Ubiquitin polyprotein (nonamer) at 1.7 M GdnHCl (A) without and (B) with the xenonucleus; and at 0 M GdnHCl (D) without and (E) with the xenonucleus; (C) Scattered plot of force vs ΔL_c_ for only ubiquitin (black) and ubiquitin with xenonucleus (red), at 1.7 M GdnHCl respectively; (F) Scattered plot of force vs ΔL_c_ for only ubiquitin (black) and ubiquitin with xenonucleus (red) when no GdnHCl is added (at 0 M GdnHCl).

### Understanding the effect of sequence specificity of the ‘xenonucleus’ using coarse-grained structure-based models

Symmetrized structure based potentials (symSBMs)^59–61^ investigate domain-swapping by simulating two identical protein chains held together by a weak constraint. These proteins are allowed to form contacts present in the folded state either within the chain or between chains. We first use a modified symSBM coarse-grained to a Cα-level to simulate the ubiquitin-peptide interaction. All simulations are performed at a temperature where both the folded and unfolded states of ubiquitin are equally populated and the transition regions are well sampled. This is similar to performing experiments at 3 M GdnHCl. We compare the kinetics and thermodynamics of three sets of simulations for every peptide construct: (1) ubiquitin without the peptide, (2) ubiquitin with the peptide and (3) ubiquitin with a pre-folded or stapled peptide in which the intra-peptide native contacts are strengthened.

When the xenonucleus peptide is present, ubiquitin can either use its own β1-β2 as a nucleus or the externally supplied one. The free energy profile of ubiquitin without the peptide is similar to that of ubiquitin with the unstapled peptide (Fig. 5A) and has only two minima, unfolded (U) and monomer (M), indicating that the protein does not prefer the unstapled peptide as a folding nucleus. However, an intermediate M’ is populated when ubiquitin is simulated with the stapled peptide. In order to understand the nature of the M’ ensemble, 2D-free energy profiles (2DFEPs) of the simulations were plotted (Fig. 5C and 5E) with the two coordinates being the number of formed contacts between ubiquitin and the peptide (Q_inter_) and the number of contacts formed within the ubiquitin chain. A significant amount of Q_inter_ are formed in M’ and further analysis of the structures populated in this ensemble (Figure S18A and S18B) indicate that ubiquitin uses the pre-folded xenonucleus instead of its own β1-β2 in M’. The ubiquitin β1-β2 also folds but interacts only loosely with the rest of the protein. The M’ minimum is also populated in the ubiquitin-unstapled peptide simulation but to a much lesser extent. An analysis of the ubiquitin-stapled peptide simulation trajectories beginning from the M’ ensemble (Table S1) shows that the protein is 5 times more likely to get unfolded (visit U) than it is to fold directly to the monomer (M). The population percentages of the trajectories imply that for M’ to reach M, it has to unfold first.

**Figure 5.**
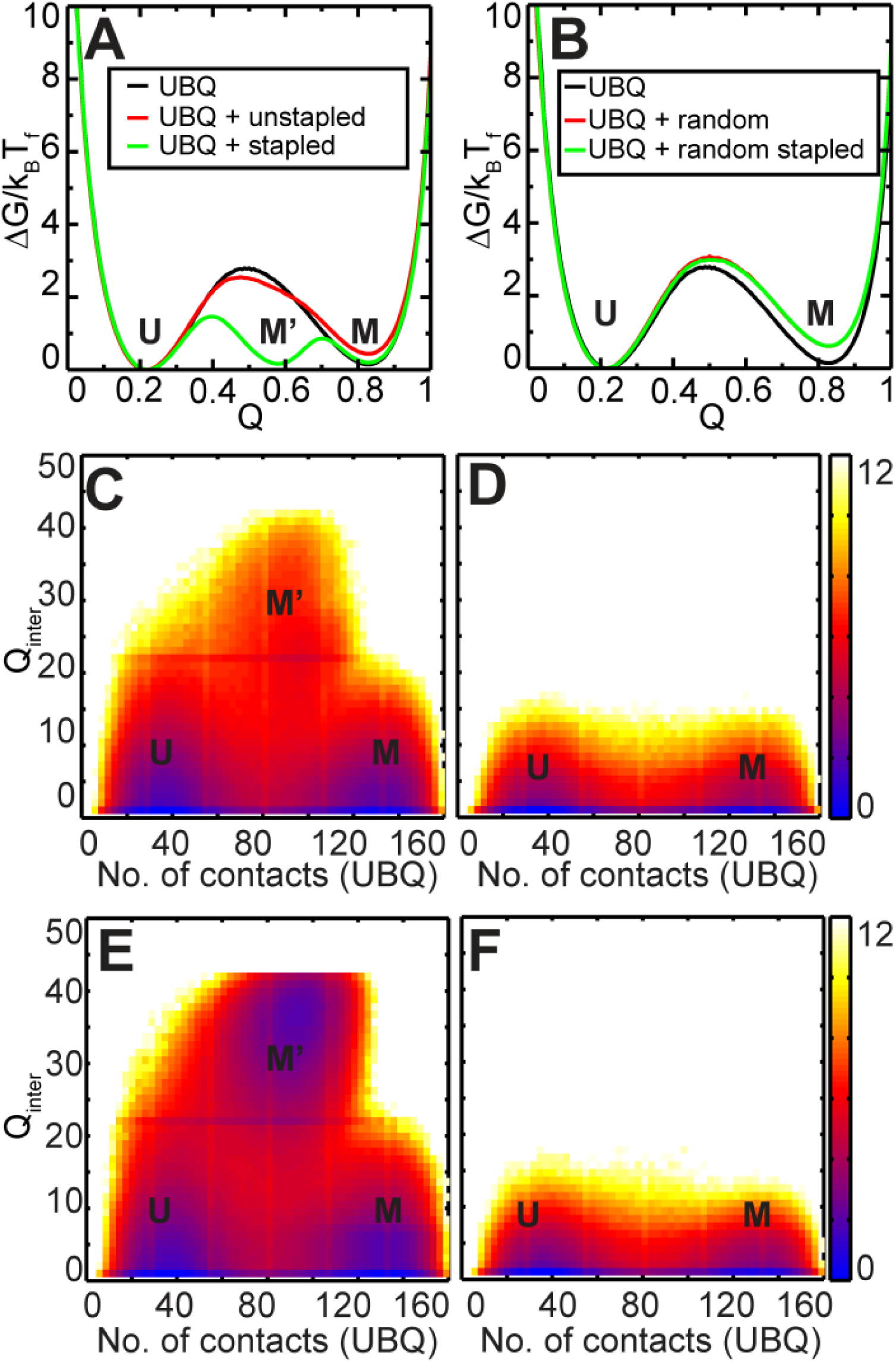
Free energy profiles for simulations of UBQ with xenonucleus (A) Free energy profile for only UBQ (black), UBQ with unstapled (red) and stapled (green) xenonucleus. C and E, 2DFEPs of Q_inter_ vs. No. of contacts (UBQ) for UBQ + unstapled and stapled xenonucleus respectively. (B) Free energy profile for only UBQ (black), UBQ with randomised contacts (both intra and inter) for unstapled (red) and stapled (green) xenonucleus with randomised contacts. D and F, 2D FEPs for UBQ + randomised unstapled and stapled xenonucleus respectively. The colours indicate the population of each minimum, with a darker colour corresponding to a higher population. Colour bars for 2D FEPs are displayed on the right.

In order to understand the effect of sequence perturbations on the action of the peptide, we simulated ubiquitin in the presence of a stapled and unstapled peptide which has the same folded structure as the xenonucleus but whose contacts with the rest of the protein are randomized. This implies that the protein-peptide contacts are not in harmony with the peptide structure and only a subset of protein peptide contacts can be made at a given time. This construct mimics the random peptide used in the experiments. The free energy profiles of these constructs (Fig. 5B) are similar to those of only ubiquitin and have only two ensembles, U and M. 2DFEPs (Fig. 5D and F) confirm this observation and there is no accumulation of the M’ state.

## Discussion

Kinetics and thermodynamics of protein folding are routinely modulated by changing solvent conditions, e. g. by the addition of urea or guanidine hydrochloride, but such changes are not protein specific. There are few examples where a specific ligand is able to bind and alter the folding of a natively folded protein^62,63^. Here we show a general approach for rationally designing such ligands by mimicking the folding nucleus of a protein. We show that the β1-β2 part (residues 1 to 17) of ubiquitin, which is known to be the segment which folds first (into a β-hairpin shape), can make folding faster when introduced as a separate conformationally constrained peptide at excess concentrations [Figure 1 (A) and (B)]. We constrain the ‘xenonucleus’ so that it can easily adopt the native-like β-hairpin shape, by engineering a disulphide bridge between the termini. Refolding kinetics, measured by using stopped-flow experiments, show that the xenonucleus can force the protein to fold faster in a concentration-dependent manner [Figure 1 (C)]. Substantial increase in the rate of the folding is observed even at a very high GdnHCl (e. g. 3 M GdnHCl) concentration. This increase is dose dependent, up to a concentration of 60 μM. This shows that the xenonucleus is the causative agent for this increase in the folding rate. The slight decrease at higher xenonucleus concentrations may be due to the partial aggregation (e.g. formation of small oligomers) of the externally added xenonucleus. We note that the xenonucleus is not covalently linked, so there is always a competition between the proteins native nucleus and the xenonucleus in interacting with the rest of the protein. The effective concentration of the native β1-β2 is of course very high, and so the effect of the xenonucleus is expected at relatively high concentrations, as has been observed. Substantially larger effects are observed in experiments which have covalently modified native folding contacts (no xenonucleus)^47^.

A weaker but still significant effect is observed when this external peptide is not conformationally restrained or is in a randomized sequence [Figure 1 (D)]. A peptide with similar secondary structure but the randomized sequence will not have the same amino acids at the corresponding positions and hence will not fit properly in the right pocket (due to different side-chain interactions). However, we may expect some non-specific interactions of an unstable protein (at high GdnHCl concentrations) with a peptide whose average properties (e.g. hydrophobicity and charge) resemble a part of the protein itself. So our results suggest that the xenonucleus increases the rate of folding of the protein through a combination of specific and non-specific interactions. We note that the external peptides are added at two hundred μM concentrations (except as noted), which is about mg/ml. Chances of any crowding effect are very low at such concentrations so the background effect is probably due to non-specific interactions.

Interestingly, the thermodynamic stability of the protein remains almost unchanged as seen in the steady-state chemical denaturation experiments (using both the Tryptophan steady-state fluorescence and circular dichroism, Figure 2 (A) and Figure 2(A) inset). Though the protein to peptide ratio used in the CD experiment could not be kept the same as that in the kinetic experiments due to the lower sensitivity of the technique, the results still indicate that the thermodynamic stability of the protein is not altered by the presence of the peptide. We infer that the xenonucleus reduces the free energy barrier for the folding-unfolding transitions, without changing the relative stability. In that sense, the xenonucleus acts as a folding catalyst.

The steady state experiments also imply that the unfolding rate must become faster by an equal amount, and we test that by measuring the unfolding kinetics. Our experiments do show that the unfolding rate becomes faster [Figure 2 (B) and (C)]. However, the increase in the unfolding rate is somewhat larger than the increase in the folding rate. We note that the folding and the unfolding experiments are not exact reverses of each other. The ensemble of unfolded states at 4.8 M GdnHCl is likely to be very different from the unfolded ensemble at 1 M GdnHCl. Therefore kinetic and thermodynamic results are not necessarily inconsistent. This result further supports our hypothesis that the externally-supplied xenonucleus reduces the energy barrier separating the unfolded and the folded states, which in turn makes both the folding and the unfolding faster without changing the ground-state energies (described in the SI, Figure S19). It can also enhance the rate by minimizing other ‘kinetic traps’ that a protein has to encounter during folding or unfolding events.

While these experiments prove that the presence of the xenonucleus in the solution causes a change of the folding kinetics, they do not prove that it does so by interacting directly with the protein. Time resolved FRET experiments provide evidence of direct molecular interaction. Our results show that the peptide directly interacts with the protein during folding [Figure 2 (D)]. We observe a clear reduction in the lifetime of Rh110 (∼15%) in the presence of the Cy5-labelled xenonucleus. The two components of the fluorescence lifetime possibly indicate that there are two conformations of the protein present in the solution. It is interesting to speculate that these are related to their folding states, but that cannot be proven from our experiments. The population of the shorter lifetime component increases strongly upon exposure to the xenonucleus, while its lifetime goes down by a factor of two. This shows that the xenonucleus is directly interacting with the protein.

The presence of such interaction still does not prove the original hypothesis, that the xenonucleus can substitute the native nucleus (i.e. the β1-β2 part) of the protein. Direct evidence for this comes from solution-state NMR experiments. It is apparent from the NMR data (chemical shift perturbations) that the xenonucleus interacts with the rest of the ubiquitin protein primarily at the sites which interface with the β1-β2 in the wild-type folded protein [Figure 3]. Interestingly, such an interaction is possible only when the protein is perturbed by adding GdnHCl (1.7 M). No such interaction occurs when the same experiment is performed in the folding buffer (0 M GdnHCl). While the perturbation is weak, suggesting that there is considerable spread in the exact mode of interaction, the location of the perturbations establish that a residue-by-residue substitution of the native β1-β2 by the xenonucleus is the preferred mode of interaction.

This specific interaction is also observed in the coarse-grained simulations of ubiquitin folding [Figure 5]. Potential energy functions which encode the structure of the protein have successfully been used in conjunction with molecular dynamics to simulate both protein folding and 3-D domain-swapping^61,64,65^. In domain-swapping, monomeric proteins exchange a portion of their structures to give dimers. The monomer structure of each of the chains in encoded by using native bond lengths, angles, dihedrals and native contacts. Additionally, any native contact that is present in between two residues within a chain can also form between the corresponding residues present in two different chains. So this provides an appropriate technique for our exploration of the interaction of the xenonucleus with the protein. When the constrained xenonucleus (5x and 10x strengths of β1-β2) is presented to the thermally unfolded ubiquitin, it can interact with the parent protein and create a nearly-folded state. We see the formation of a new intermediate state (M’), which is seen as an extra dip in the free energy plot apart from U and N. This intermediate state corresponds to the ubiquitin folded on the xenonucleus instead of its β1-β2. Overall, M’ buffers the protein, acting like a holdase^66^. In contrast, the unconstrained peptide or the randomized sequence can hardly do the same. We note that this scenario may slow down folding, and not speed it up as we have observed here. However, if the final state we observed in our stop-flow measurements is the xenonucleus attached state (M’), then the process from U to M’ may become faster than the actual folding kinetics. This is possibly the end-state we observe in our kinetics experiments. Indeed, our FRET results, which are performed in the steady state at 3 M GdnHCl, show that the peptide remains attached to the folded protein [Figure 2 (D)]. These results show that a stably folded peptide which can make a substantial number of contacts with the protein can act as a xenonucleus.

Further proof of steady state interactions comes from the single-molecule force unfolding experiments. It also helps answer the question about the fate of the β1-β2 part when the xenonucleus occupies its position in the protein. If this part is free, then it should be able to act as the ‘xenonucleus’ for a neighboring copy of the protein, producing a dimer. The local concentration of the protein must be high for such an interaction, and ubiquitin nonamers provide a model for this. We observe that the unfolding of ubiquitin nonamers under slightly destabilizing conditions (1.7 M GdnHCl) is strongly affected by the xenonucleus. Insertion of xenonucleus as β1-β2 of ubiquitin changes its contour length, as the original β1-β2 of the protein is free [Figure 4]. The contour length is occasionally double of what is expected. Furthermore, the mechanical stability of the xenonucleus-inserted ubiquitin is different as the pulling geometry is now from β3-C-terminus. Interestingly, no xenonucleus effect is observed when the protein is mixed with the peptide at 0 M GdnHCl. It indicates that the xenonucleus indeed interacts with the protein at 1.7 M GdnHCl and gets incorporated into the folded structure. Further, neighbouring proteins get interlinked so that they unfold as a unit, increasing the contour length. This is consistent with the interpretation that the free β1-β2 part of one protein can insert itself into the neighbouring protein, a scenario with similarity to 3-D domain swapping^61,64,65^. This happens most often at 1.7 M GdnHCl and only rarely at 0 M GdnHCl, likely because the protein is fluctuating between the folded and the unfolded forms more often at the higher GdnHCl concentration.

Our results suggest that the folding of almost any protein that possesses a well-defined folding nucleus can be modulated by introducing a nucleus-mimicking molecule. Since this is a peptide sequence, in principle, it is possible to introduce such a nucleus inside a living cell, either through an external route or by simply expressing it. We expect proteins to be mostly folded inside the cell, and perhaps the xenonucleus will not affect its target protein, except during the process of translation. However, misfolding is common inside the cells^9,67–69^, and a protein in the cell can unfold and fold several times during its lifetime. A change in these rates, induced by the xenonucleus, can significantly alter the cellular processes involving the protein. Depending on how significant or ubiquitous these processes are, the cell physiology itself may be affected by the xenonucleus. It also implies that fragments of proteins that are naturally present^70–76^ as degradation by-products *in vivo* can, in principle, affect the folding of the full-length proteins^74,77,78^. It is likely that this chaperone will be rather specific for the particular protein, or at least for a class of homologous proteins that it has been designed for. The nuclei in general may have a family of common shapes, such as the beta hairpin shape appropriate for ubiquitin. To that extent, other proteins that share a common theme for the folding nucleus, without much sequence homology, may also get affected by the decoy nucleus. For example, in designing small molecules or peptides which would interfere with the formation of protein complexes by attaching to the protein–protein interface^79–81^, it has been observed that multiple mimics with secondary structure similarities can eventually do so at a similar extent. In any case, this short peptide may be expected to act as a micro-chaperone for a small subset of proteins. We note that the xenonuclei may also be constructed from non-natural peptide mimics, which can be potentially used in pharmaceutical formulations in vitro. If such molecules are cell-permeable, then it can also provide a potential new route for selective chemical intervention in the folding of cellular proteins.

## Cartoon

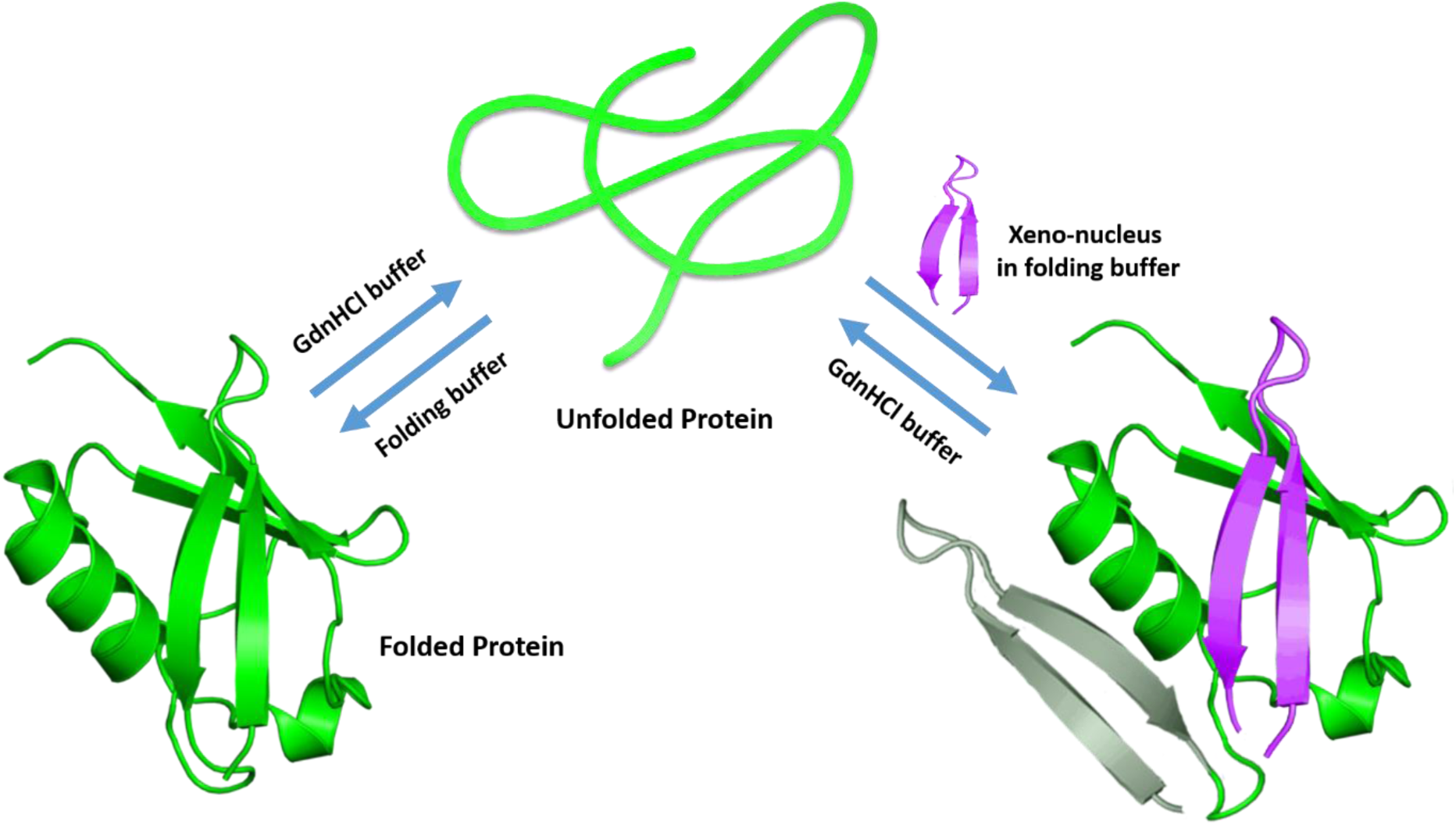

### Experimental Section

## Methods

### 1. Synthesis of L-peptides and formation of disulphide bond

All the peptides used in this work are synthesized on Rink Amide MBHA resin using 9-fluorenylmethoxycarbonyl (Fmoc) chemistry in an automated peptide synthesizer (PS3, Protein Technologies, USA). Disulphide bonds are formed by I_2_-oxidation of acetaminomethyl (Acm)–protected cysteine residues and we have named these disulphide-bridged peptides as ‘stapled peptides’. The peptides are subsequently purified by reverse-phase high-performance liquid chromatography (HPLC) and characterized in the laboratory following a well-established protocol described previously^82^.. The sequence of the stapled Cys-Ubq3-Cys peptide used as the xenonucleus is chosen from WT Ubiquitin itself; **C**-MQIFVKTLTGKTITLEV-**C** (the middle part consists the same 1 to 17 residues present in wild-type (WT) Ubiquitin), and the Cys-Ubq3 is only slightly different from this for not having the C-terminal Cysteine residue (**C**-MQIFVKTLTGKTITLEV). Truncated versions of Cys-Ubq3, namely Cys-Ubq1 (**C**-MQIFVKTL) and Ac-Ubq2 (**Ac**-GKTITLEV), were synthesized separately. Here, Ubq1 and Ubq2 represent the β1 and β2 strands present in the wild-type ubiquitin. In Ac-Ubq2, free amine at the N-terminal has been acetylated to mimic the peptide bond as closely as possible. For the Cys-Random Ubq3-Cys (the ‘Stapled Random’ peptide) and Random Cys-Ubq3 (the ‘Random’ peptide) peptides, the same residues as those in the β1-β2 (MQIFVKTLTGKTITLEV) of WT ubiquitin are completely randomized, and the sequences are namely, **C**-TGVTMVFIELLQKTTKI-**C** and TGVT**C**MVFIELLQKTTKI. The formation of the disulphide-bridge is characterized by checking the mass using Matrix Assisted Laser Desorption/Ionization (MALDI-TOF) Mass Spectrometry and Electrospray Ionization Mass Spectrometry (ESI-MS), described in the SI, Figure S2-S7. For the Cyanine5-labelled stapled Cys-Ubq3-Cys, the dye (Cy5, Cyanine5 NHS ester, Lumiprobe) was covalently linked to the free amine at the N-terminal while on beads, similarly cleaved and purified subsequently (mass spectrum shown in the SI, Figure S8).

### 2. Synthesis of the D-peptide segment (Stapled D-peptide)

The D-ubiquitin peptide segment cmqifvktltGktitlevc was synthesized using an automated peptide synthesizer (Tribute-UV/IR from Protein Technologies, USA). Fmoc-SPPS was carried out in 0.1 mmol 2-Cl-(Trt)-resin using amino acids (AA) (2.5 equiv., 0.25 M), DIC (0.25 M) as a coupling reagent and Oxyma (0.25 M) with DIEA (0.025 M) as additives. Cysteine residues were coupled for 2 min at room temperature followed by 5 min at 50 °C. All other amino acid couplings were performed for 7 min at 50 °C under N_2_ atmosphere with vortex mixing. After the first residue coupling on the resin, the unreacted functional group on 2-Cl-(Trt)-resin was capped using 5% MeOH (vol/vol) in DMF. Fmoc deprotection after every coupling cycle was carried out by 20% piperidine treatment at 50 °C. After synthesis, the D-peptide was cleaved from the resin using TFA (85%), Phenol (5%), TIPS (2.5%), Water (5%) and DODT (2.5%) as a cleavage cocktail. After cleavage, the TFA was evaporated under N_2_ flow inside a well-ventilated fume hood. The cleaved peptide was precipitated and washed with diethyl ether. After lyophilization, 30 mg (33 µmol, one-third of the total crude peptide) of the dry crude peptide was dissolved in 10% aqueous DMSO (30 mL) and incubated at room temperature for 24 h to furnish the oxidized (disulfide bonded) D-peptide. The crude oxidized D-peptide was then loaded directly on to a preparative HPLC column for purification, which afforded 3 mg (1.4 µmol, 4% yield) of the pure disulfide stapled D-peptide (Figure S9). Observed mass (ESI-MS): 2126.11 ± 0.01 Da (deconvoluted most abundant isotope); calculated mass: 2126.62 Da (average isotope).

### 3. UbqF45W expression and purification

WT ubiquitin cloned in pQE80l vector is mutated to UbqF45W using site-directed mutagenesis and the clone is confirmed positive using Sanger’s sequencing method^83,84^. The plasmid containing the UbqF45W gene is then transformed into E.Coli BL21 (DE3) cells and induced with 1mM IPTG at 0.6 OD_600_ for 6 hours. The cell pellet obtained is resuspended in 1X protease inhibitor cocktail containing 20 mM sodium phosphate-buffered saline (20 mM PBS, pH 7.4) and then lysed by sonication at 4°C. The lysate obtained after centrifugation at 17000 rpm for 45 min at 4°C is incubated with Ni-NTA resins for 4 hours at 4°C on rotaspin. Ni-NTA resins incubated with lysate are loaded in a 15 ml chromatography column. After collecting flow through beads are washed with 20 mM PBS (pH 7.4) and 20 mM PBS (pH 7.4) containing 20 mM imidazole to get rid of non-specific impurities. The desired protein is then eluted using 20 mM PBS (pH 7.4) containing 250 mM imidazole. The protein is further purified by size exclusion chromatography using a Superdex 75 column on the Bio-Rad Biologic Duo-Flow FPLC system and characterized using MALDI-TOF mass spectrometry (Figure S1).

### 4. Polyprotein expression and purification

BL21 DE3 strain of *Escherichia coli* cells are transformed with pQE80L-(ubq)_9_ construct. The cells are grown in LB medium in the presence of ampicillin (0.1 mg/ml) at 37°C and 200 rpm till OD_600_ reaches ∼0.6. The polyprotein is overexpressed by IPTG induction (1mM) and the culture is allowed to grow for ∼6 hours. The cells are subsequently harvested and lysed by sonication using lysis buffer (containing leupeptin, pepstatin, lysozyme, PMSF, and Triton-X). The expressed protein is first purified by affinity chromatography using Ni-NTA coated agarose beads (Roche). His_6_-tagged proteins are eluted using 250 mM imidazole. The polyprotein is further purified using size-exclusion chromatography using the Superdex200 column (Amersham Biosciences). The purity of (ubiquitin)_9_ is checked using sodium dodecyl sulphate polyacrylamide gel electrophoresis (SDS-PAGE).

### 5. Refolding experiment by Stopped-Flow Fluorescence

The stopped-flow fluorescence measurements have been carried out on a MOS-250 Spectrometer equipped with a stopped-flow accessory (Biologic, SFM-300). The time-resolution of the stopped-flow instrument has previously been measured by Nandi et al.^85^, by using the standard NATA-NBS quenching experiment. The kinetic rate constant obtained by them (8.5 × 10^5^ mol^−1^ second^−1^) was in good agreement with the previously reported values in the literature^86,87^. As WT Ubiquitin does not have any intrinsic fluorophores, we have chosen the F45W mutant. Its biophysical properties have already been well studied^57,88–90^. In a typical refolding experiment using the 3-syringe stopped-flow fluorescence, the protein Ubiquitin F45W is first unfolded at 4.25 M Guanidine hydrochloride (GdnHCl) and then rapidly diluted with the saline phosphate buffer solution (PBS, Composition: 146 mM NaCl, 5.4 mM KCl, 0.4 mM KH_2_PO_4_, 20 mM Na_2_HPO_4_ with 2 mM NaN_3_) through a mixer (total volume = 251 µL and flow rate = 10 mL/s). The flow is then suddenly stopped and the reaction inside the optical cell is monitored by using a PMT detector (described in the SI, Figure S9). The acquisition is started 10 ms before the stop. In a 3-syringe stopped-flow experiment, the three different syringes contain the folding buffer 20 mM PBS at pH 7.5 (Syringe 1, S1), 4.25 M GdnHCl (Syringe 2, S2), and 30 µM unfolded protein in 4.25 M GdnHCl (Syringe 3, S3), respectively. The volume ratio of S1 and S2 are changed to create jumps to different final GdnHCl concentrations, such as 1.7 M, 2.1 M and 3 M, keeping contents of S3 the same (achieving 3 µM final concentration of the protein in all the cases). The content of the syringe 1 is replaced by the peptide solution in PBS (at 200 µM, for all except for the cleaved Cys-Ubq3 fragments, which were added at 115 µM due to the lower solubility of the Cys-Ubq1 fragment) for the refolding experiments in the presence of the xeno-peptides. The peptide stocks were prepared by dissolving the lyophilized peptides in 20 mM PBS. The solution was vortexed for approximately 10-15 minutes, and any aggregates were separated by centrifugation (at 2000 × *g* for 30 minutes). The concentration of the supernatant was checked by its absorbance at 258 nm on an Analytikjena, SPECORD 205 UV-Vis spectrophotometer. The dissolved peptide was then flash-frozen in liquid nitrogen and stored at -80°C. Stocks were thawed quickly before use. The stopped-flow instrument recorded the changes in the fluorescence intensity as a function of time (excitation, 295 nm; collection at > 320 nm using a cut-off filter). All the experiments were performed at room temperature (25°C).

### 6. Steady-state denaturation by tryptophan fluorescence

Steady-state fluorescence emission spectra are recorded on a Fluorolog 3, Horiba, Jobin Yvon Spex spectrofluorimeter. 50 µL of the folded Ubiquitin F45W protein (100 µM) is added to a 450 µL mixture of PBS and GdnHCl, the ratio of the latter two are changed to increase the final GdnHCl concentration from 0 M to 5.3 M. In the second case (with the stapled xeno-peptide), 100 µL of PBS is replaced by the peptide solution in PBS (200 µM stock concentration) at each GdnHCl concentrations. The concentration of the protein and the stapled peptide in all the GdnHCl are thus kept constant at 10 µM and 40 µM, respectively. Fluorescence spectra of each of these solutions (different GdnHCl concentrations) are recorded by exciting at 295 nm (tryptophan) and collecting the emission from 310 nm to 450 nm, with an increment of 1 nm and an integration time of 0.10 s (excitation and emission slits were kept at 5 nm/5 nm, respectively). The experiments were conducted at room temperature (25°C).

### 7. Steady-state denaturation by far-UV circular dichroism

Steady-state far-UV circular dichroism spectra are recorded on a J-1500, JASCO Circular Dichroism Spectrophotometer. 25 µL of the folded Ubiquitin F45W protein (200 µM) is added to a 475 µL mixture of PBS and GdnHCl, the ratio of the latter two are changed to increase the final GdnHCl concentration from 0 M to 5.6 M. In the other case (with the stapled xeno-peptide), 135 µL of PBS is replaced by the peptide solution in PBS (148 µM stock concentration) at each GdnHCl concentrations. The concentration of the protein and the stapled peptide in all the GdnHCl are thus kept constant at 10 µM and 40 µM, respectively. Far-UV circular dichroism spectra of each of these solutions (different GdnHCl concentrations) are recorded by monitoring the signal (absorbance) from 260 nm to 200 nm with a data pitch of 0.5 nm, CD scale of 200 mdeg/1.0 dOD, data integration time of 1 sec, Bandwidth of 2.00 nm and average of 3 accumulations.

### 8. Unfolding kinetics of ubiquitin by Stopped-flow Fluorescence

Unfolding kinetics of ubiquitin is measured using the same 3-syringe stopped-flow fluorescence set-up as described earlier. Ubiquitin F45W (30 µM) is pre-mixed and equilibrated at 1 M GdnHCl before starting the kinetics experiment (Syringe 3, S3). Then a jump is created to a final GdnHCl concentration of 3 M by rapidly mixing with 4.8 M GdnHCl (Syringe 1, S1) and 20 mM PBS (pH 7.5) with azide (Syringe 2, S2) in the ratio of S1:S2:S3 = 6:3:1. In the other case, unfolding of ubiquitin in the presence of the stapled Cys-Ubq3-Cys, Syringe 2 solution is replaced by the stapled peptide in pH 7.5 20 mM PBS with azide, keeping others the same. Total flow volume was kept at 251 µl and the total flow rate 10 mL/s, the acquisition was started 10 ms before the stop.

### 9. Fluorescence lifetime experiments

The fluorescence lifetime experiments were performed in a commercial confocal microscope based setup (LSM-710, Carl Zeiss). 486 ± 14 nm laser light was obtained from the OYSL supercontinuum white-light laser source with the help of home-built spectrometer made up of two prisms and a slit. The repetition rate of the laser pulses was kept at 5 MHz. The light was guided to the sample through the IR laser path of LSM. Excitation light reaches the sample after passing an 80/20 beam splitter which allows 20% excitation light to pass and a mirror right below the 40X water immersion objective lens of 1.2 NA. The fluorescence photons were collected using the same objective lens via descanned path of LSM. After filtering the excitation light using a suitable emission filter, a 5 cm convex lens was used to focus the fluorescence on a fiber of core diameter 100 µm. Another side of fiber was coupled to MPD detector (Micro Photon Devices) which sent the signal to a time-correlated single-photon counting (TCSPC) card SPC150 (Becker & Hickl GmbH). For all the lifetime measurements, data were recorded until the maximum count becomes greater than 10,000.

### 10. Single-molecule force spectroscopy (SMFS)

All SMFS pulling experiments are carried out on a custom-built atomic force microscope described elsewhere^91^. In all SMFS experiments, about 60 μL sample (prepared in 20 mM PBS, pH 7.4) is added onto a gold-coated coverslip and allowed to rest for ∼20 minutes. We use gold-coated reflective cantilevers with silicon nitride tips from Bruker, CA, USA. Spring constant of the cantilevers is measured using the equipartition theorem as previously described. The spring constant of the cantilevers that we use varies in the range ∼40-70 pN/nm. All force-extension experiments are carried out at ambient temperatures. The pulling speed in all SMFS experiments is 400 nm/s. Ubiquitin nonamer (∼140 μM) is used. GdHCl concentration is maintained at 1.7 M in all experiments to see the effect of GdHCl on the mechanical stability of ubiquitin. A concentration ratio of 1:1 (stapled peptide: ubiquitin (monomeric concentration)) is maintained to see the effect of the stapled peptide on the mechanical properties of ubiquitin. GdHCl and stapled peptide in all samples are incubated with (ubiquitin)_9_ for ∼1 hour at 4°C to allow sufficient equilibration before starting AFM experiments.

### 11. SMFS data analysis

Force vs Extension (FX) traces containing even a single protein unfolding peak is considered for data analysis. These FX traces are fitted using the worm-like chain (WLC) model of polymer elasticity using equation 1^92,93^.

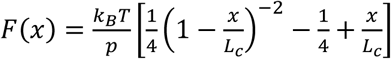

where F, x, p, L_c_, k_B_, and T denote the force, the end-to-end molecular extension, the persistence length, the contour length, the Boltzmann constant, and the absolute temperature, respectively. The persistence length values used for fitting varies between 0.13 to 0.84 nm.

### 12. Ubiquitin (wild-type) expression and purification for NMR experiments

For protein purification, the sequenced clone was transformed in BL21 (DE3) bacterial cells and plated. A single colony was selected to grow in 10 ml Luria Bertani (LB) broth for 6 hours. Cells were pelleted and transferred in 50 ml M9 minimal media with ^13^C Glucose and ^15^N Ammonium chloride. After overnight incubation at 37°C, the grown culture was scaled to 1 liter of the same minimal media. The culture was induced at O.D. of 0.8 with 1 mM IPTG and further incubated at 37°C for 3 - 4 hours. Cells were pelleted and resuspended in 50 mM sodium acetate pH 4.5 - 4.75 and 5 mM EDTA. The theoretical isoelectric point (pI) of Ubiquitin is 6.5. Sonication at 50% amplitude for 20 cycles of 5 secs on and 25 secs off lysed the bacterial cells. The sample was then centrifuged at 17000 rpm for 45 min to separate soluble proteins from cell debris. Dilute acetic acid was added to the supernatant with continuous stirring to bring pH to 4.5 - 4.75. It was further centrifuged at about 17000 rpm (∼20000g) to remove precipitates. The supernatant was filtered and incubated with Sepharose SP (GE Healthcare) column. Bound proteins were eluted with a continuous salt gradient of 0 to 600 mM NaCl. After checking on SDS-PAGE, fractions containing Ubiquitin were pooled and further purified by Size Exclusion chromatography on Superdex 75 column (GE Healthcare) in 20 mM Phosphate Buffer Saline (PBS). Ubiquitin protein fractions were pooled, concentrated to 1 mM, and stored at 4°C until use in the presence of 0.03% wt/vol Sodium Azide.

### 13. Sample Preparation for NMR

NMR experiments were carried out in Bruker Avance III 600MHz equipped with cryo (TCI) probe. The temperature of the probe was calibrated with Methanol-*d*_4_ (99. 99%). Data processing and analysis were performed by using NMRpipe and Sparky, respectively. The sample preparation and NMR studies were performed as follows: (1) Two sets of PBS buffers were prepared for NMR studies: (a) 20 mM of Phosphate Buffer Saline (PBS), and (b) 8 M GdnHCl in 20 mM PBS. (2) 1 mM of Ubiquitin sample was diluted to 500 µM with standard 20 mM PBS. (3) The sample was equilibrated at 4.3 M GdnHCl. (4) 1 mg of L/D xenonucleus (stapled Cys-Ubq3-Cys) was dissolved in 20 mM PBS buffer. The solution was vortexed for approximately 10-15 minutes, and the soluble part was then separated by centrifugation (@2000g for 30 minutes). The dissolved peptide was stored. (5) For NMR experiments, a sample of Ubiquitin in 1.7 M GdnHCl was prepared by mixing 2:3 ratio of Ubiquitin in 4.3 M GdnHCl (step 3) and the peptide solution (step 4). (6) For the control experiment, two types of Ubiquitin samples were prepared: (a) First, Ubiquitin in 1.7 M GdnHCl was prepared by mixing a 2:3 ratio of 4.3 M GdnHCl Ubiquitin solution (step 3) and 20 mM PBS buffer. (b) For the second sample, Ubiquitin (no GdnHCl) and stapled peptide solution were mixed in 2:3 ratios. (7) A total of 275 µL (with 10% D_2_O) of the sample was loaded in 5 mm Shigemi tubes for NMR experiments.

### 14. SBM MD simulations

Structure based models (SBMs) encode the funnel shaped energy landscape of natural proteins^94^. We use Cα coarse grained SBM to simulate Ubiquitin (PDB ID: 1UBQ). Details about the energy function have been described previously^95^. We use a cut-off based contact map to simulate UBQ. This criterion defines a contact between two residues i and j if any heavy (non-hydrogen) atom of i is within 4.5 Å of any heavy atom of j. This definition results in 159 contacts for UBQ. This choice of SBM and contact map is known to correctly capture the folding pathway of UBQ^96–98^.

We use modified .gro and .top files obtained from the SMOG webserver^99^ to perform Cα SBM MD simulations using GROMACS^100^. These simulations are performed at *T*_f_, *T*_f_ being the temperature at which the protein spends equal time in folded and unfolded ensembles. SBMs use reduced units, i.e. the length scale, time scale, mass scale and energy scale are set to 1. The respective units in GROMACS (nm, ps, amu, and kJ/mol) were used except for the temperature. GROMACS uses k_B_ = 0.00831451 and calculates k_B_T as the reduced temperature. Hence, to obtain a reduced temperature of 1 GROMACS temperature needs to be 120.2717 K. All simulations were performed at 121 K, the T_f_ of UBQ. We used the leapfrog stochastic dynamics integrator with a time step of 0.0005 ps. The simulations did not include a box or periodic boundary conditions (pbc). The simulations at T_f_ were initiated using unfolded structures obtained from 1.1-1.2 T_f_. Data obtained from multiple simulations with a total of about a 100 transitions was concatenated to obtain the final dataset.

We use a symmetrized version of the Cα SBM to simulate UBQ and the xenonucleus^59,60^. Instead of two chains of UBQ, only one complete chain of UBQ is used with the other chain consisting of only the β1-β2 residues (1-14). Contacts of the UBQ chain and the β1-β2 fragment were symmetrized. UBQ and the β1-β2 fragment were held together by a weak harmonic restraint of Kb = 1 kJ/mol at a distance of r_0_ =0.5 nm. The β1-β2 fragment was simulated with intra-chain contacts scaled to strengths of 1x and 5x those of all other contacts. The 1x strength, is a mimic of the experimental condition where both UBQ and β1-β2 are allowed to refold simultaneously. Whereas at 5x, the simulations mimic the experiments where a stapled β1-β2 (xenonucleus) is added in the refolding experiments of UBQ.

Simulations of the randomized set were performed by randomizing all the contacts (both intra and inter-chain) of the β1-β2 fragment. At a strength of 1x these simulations mimic the experiments with randomized β1-β2 fragment while at the strength of 5x these simulations correspond to experiments with randomized and stapled β1-β2 fragment.

### Free energy profiles

SBMs are based on contact maps. Hence, the fraction of formed native contacts (Q) gives a good measure of how folded the protein is and is commonly used as a reaction coordinate^95^. The simulation frames with a given Q value were binned to obtain a histogram. The negative logarithm of this histogram provides the one dimensional free energy profile (FEP), i.e, a plot of the scaled free energy, ΔG/k_B_T_f_ vs. Q. The baseline of the FEPs was adjusted such that the lowest free energy point is set to 0.

Two dimensional free energy plots (2D FEPs) of Q_inter_ vs. No. of contacts (UBQ) were also plotted. Q_inter_ stands for the contacts formed between the extra fragment and UBQ (the contacts of the fragment and the same section of UBQ were not counted). No. of contacts (UBQ) stands for contacts of only the UBQ chain. Both intra and inter-chain Q values were calculated for each simulation frame and binned to obtain a 2D histogram, the negative logarithm of which gives the 2D FEPs. The baselines of these plots were adjusted such that lowest free energy value is 0.

### Trajectory analysis

Simulations at T_f_ give multiple transitions between the folded and unfolded states. We analyzed transitions from the M’ state to either U or M state. M’ is the state in which UBQ folds onto the xenonucleus instead of its own β1-β2 increasing the value of Q_inter_ while decreasing the number of intra-UBQ contacts. U and M represent the unfolded and the folded states of UBQ. A transition is counted only if the protein starting from the M’ state, visits the folded/unfolded state without visiting any other state.

