## Supporting information for "Rational design of protein-specific folding modifiers"

1.

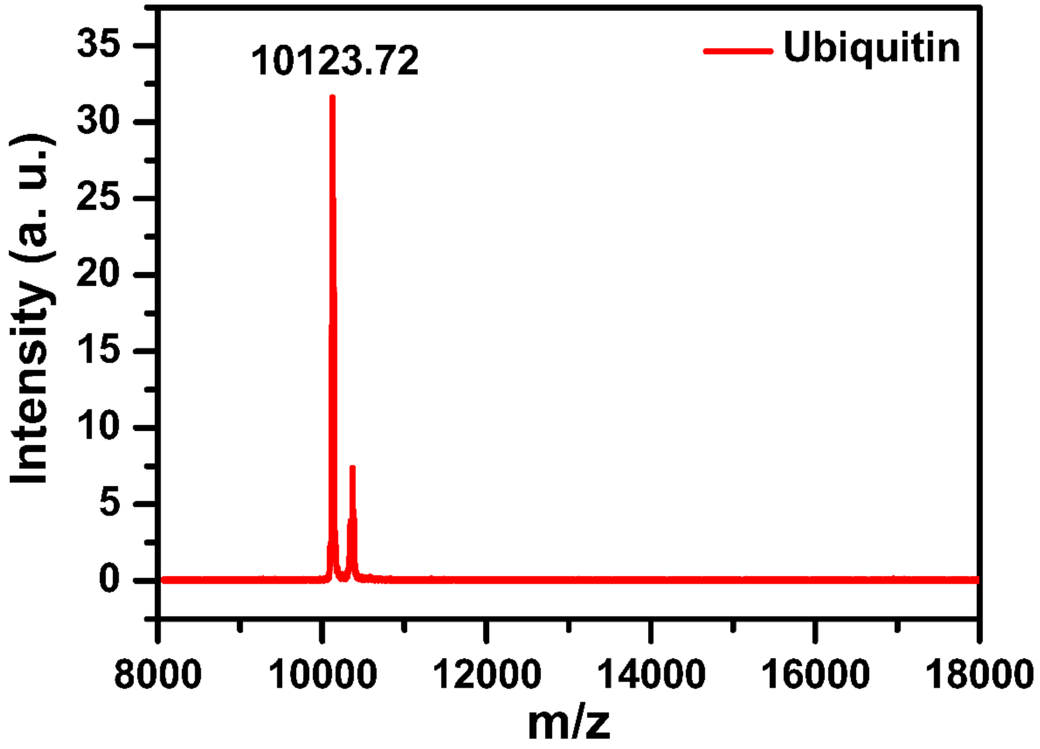


**Figure S1.** MALDI Mass Spectrum of the whole ubiquitin (UBQ) F45W protein after FPLC purification.

2.

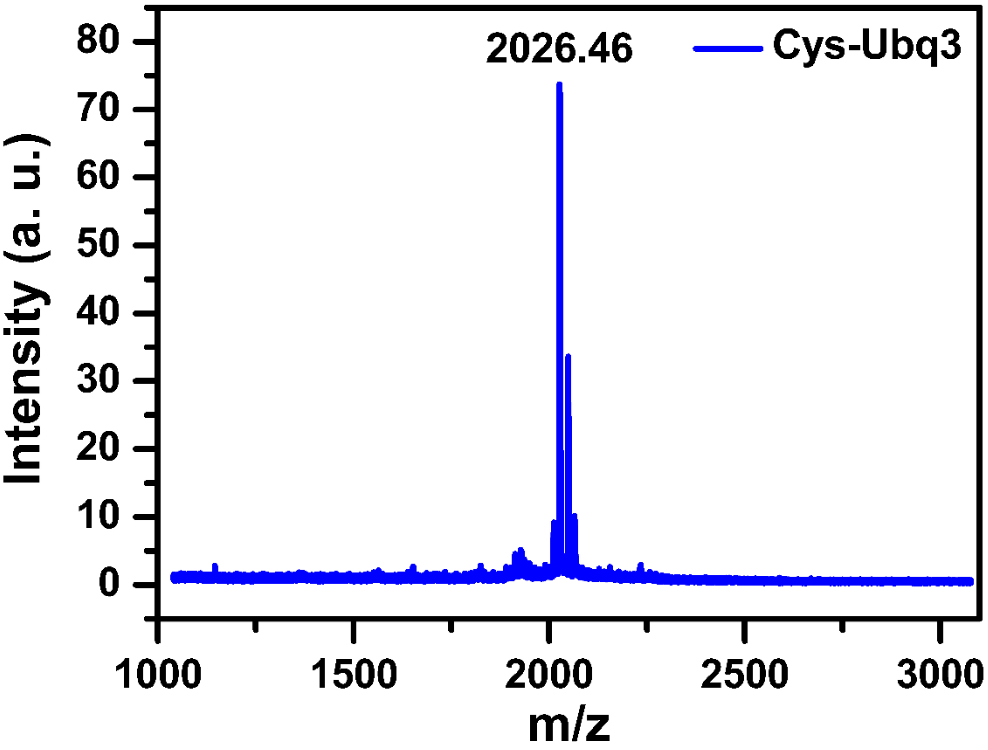


**Figure S2.** MALDI Mass Spectrum of the Cys-Ubq3 Peptide (Peptide sequence: **Cys-MQIFVKTLTGKTITLEV**).

3.

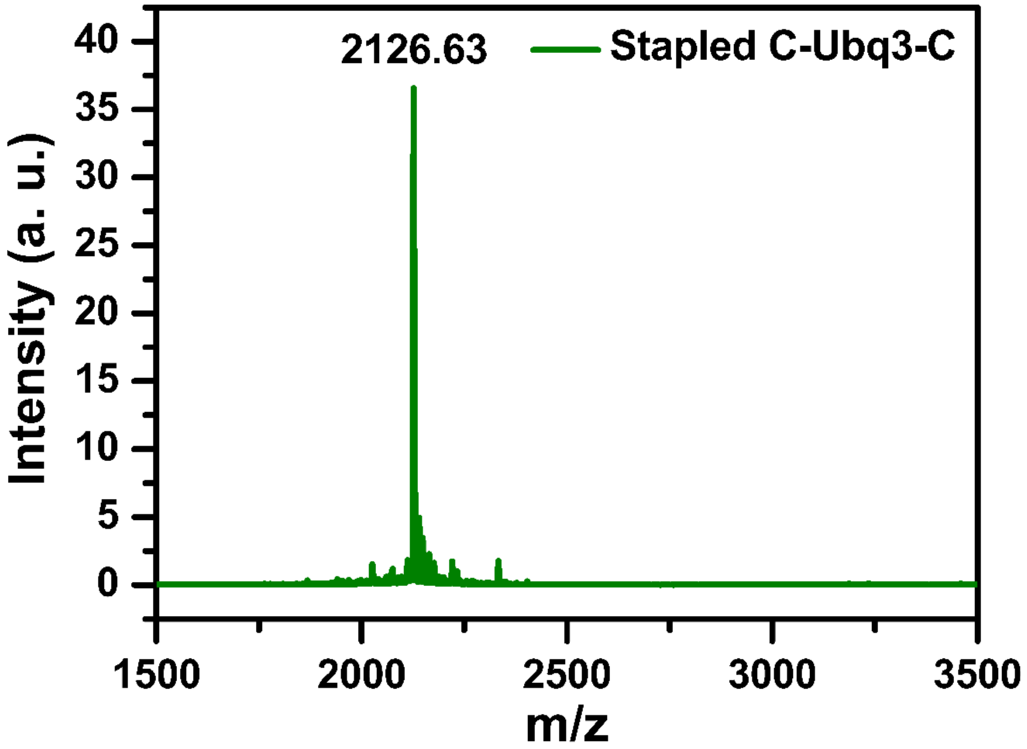


**Figure S3.** MALDI Mass Spectrum of the stapled Cys-Ubq3-Cys Peptide (Peptide sequence: **Cys-MQIFVKTLTGKTITLEV-Cys**).

### 4. ESIMS of $I_2$ Treated Stapled Cys-UBQ3-Cys in DCM, MeOH and $H_2O$:

**Chromatogram View**: Scan Segment #1

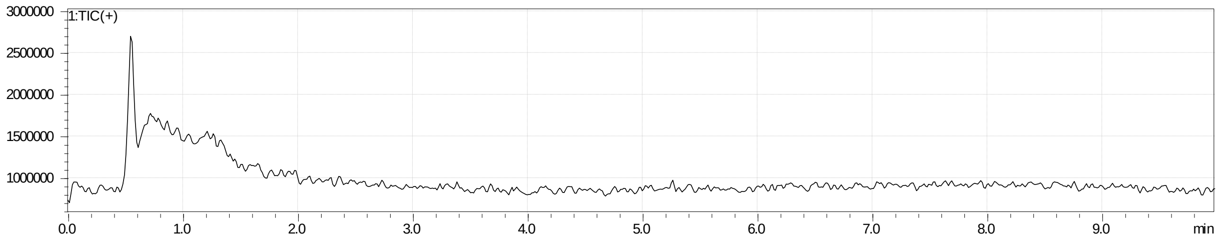


**Spectrum View:**

Event#: (1) **Scan (E+)** Ret. Time: [0.387->4.000]-[0.000->0.387] Scan#: [117->1201]-[1->117]

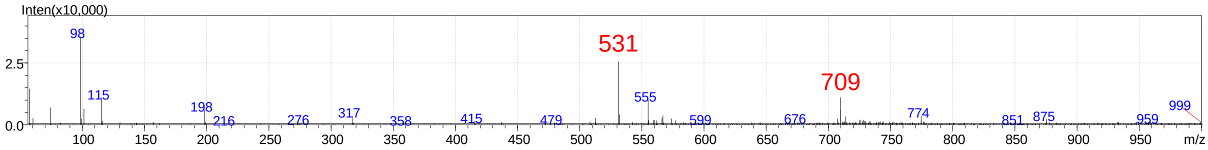


Event#: (2) **Scan (E+)** Ret. Time: [0.393->4.006]-[0.006->0.393] Scan#: [119->1203]-[3->119]

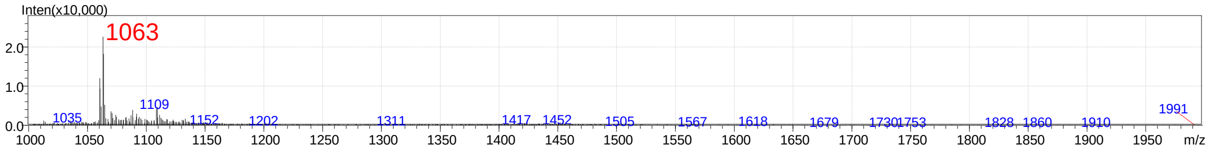


Event#: (3) **Profile (E+)** Ret. Time: [0.395->4.008]-[0.008->0.395] Scan#: [120->1204]-[4->120]

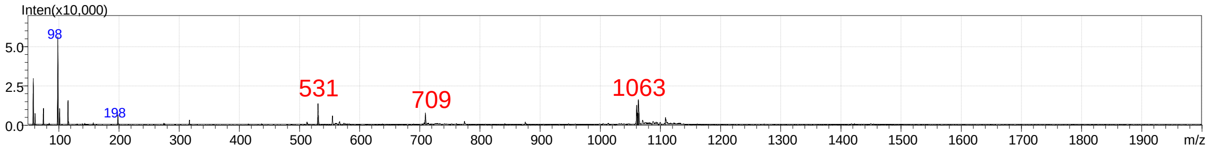


**Figure S4.** ESI-Mass Spectra (MS) of the stapled Cys-Ubq3-Cys peptide showing the presence of the disulphide bridge. The sample was dissolved in 50% Acetonitrile (ACN) in Milli-Q water with 0.1% trifluoroacetic acid (TFA) and directly injected in the MS, a constant flow (0.2 mL/min) of 50% Solvent B (0.01% Formic acid in Acetonitrile) in Solvent A (0.01% Formic acid in Water) was run through a guard column (for 10 minutes) for the analysis. The elution profile is shown in the chromatogram view (top-most panel), while the corresponding ESI-MS spectrum in the positive mode is shown in the spectrum view (below), Event#: (1) from m/z = 0 to 1000, (2) m/z = 1000 to 2000 and (3) Profile mode: from m/z = 0 to 2000, with the $[M + 2H]^{2+}$, $[M + 3H]^{3+}$ and $[M + 4H]^{4+}$ peaks highlighted in red.

5.

**(A) Mass 1063:**

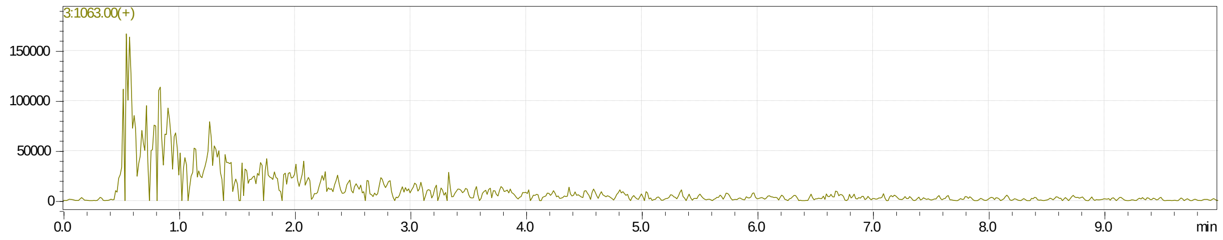


**(B) Mass 709:**

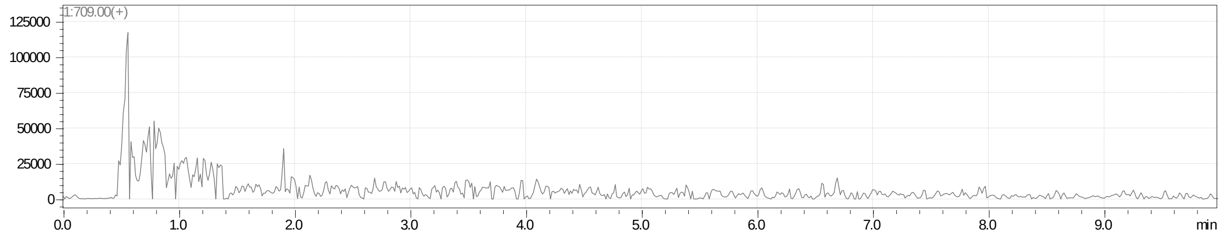


**(C) Mass 1064:**

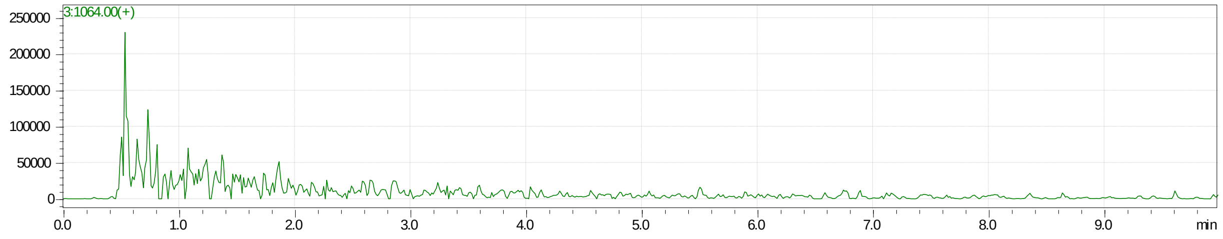


**(D) Mass 710:**

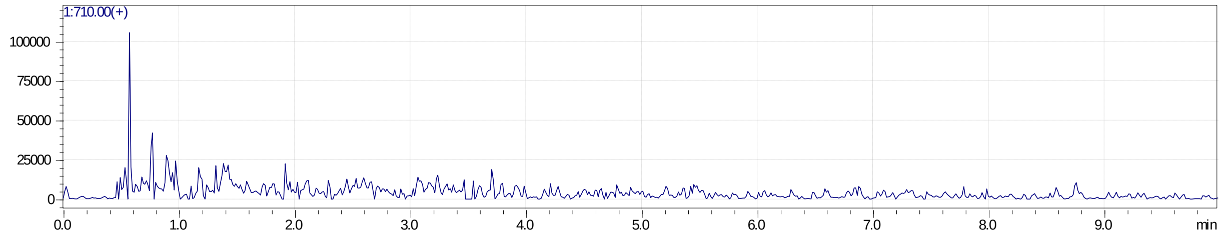


**Figure S5.** Chromatogram views (analysis of Figure S4) showing the elution of (A) $[M + 2H]^{2+}$ = 1063, (B) $[M + 3H]^{3+}$ = 709, (C) $[M + 2H]^{2+}$ = 1064 and (D) $[M + 3H]^{3+}$ = 710 peaks, clearly suggesting the presence of the disulphide bridge between the two termini of the Cys-Ubq3-Cys peptide.

6.

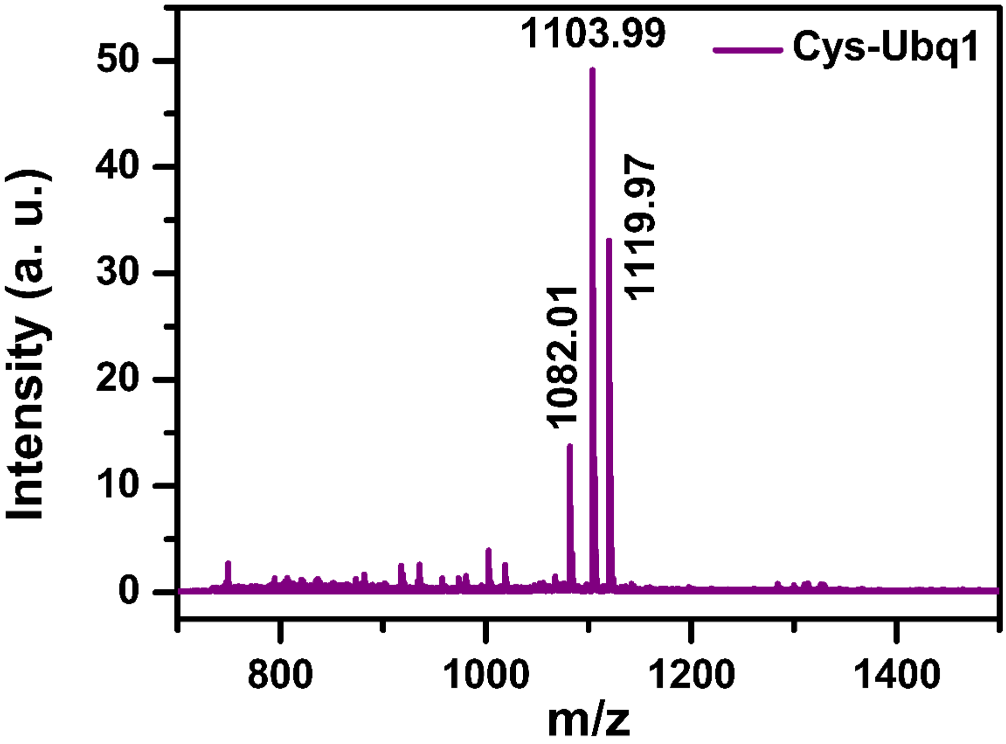


**Figure S6.** MALDI Mass Spectrum of the Cys-Ubq1 Peptide (Peptide sequence: **Cys-MQIFVKTL**).

7.

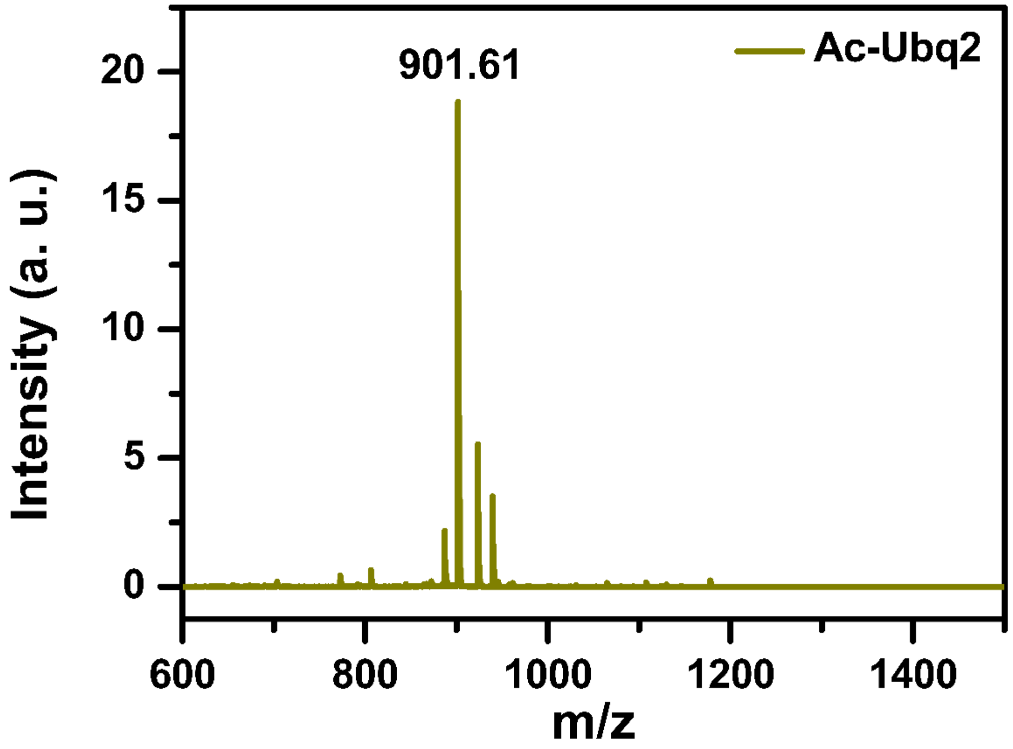


**Figure S7.** MALDI Mass Spectrum of the Ac-Ubq2 Peptide (Peptide sequence: **Ac-GKTITLEV**).

8.

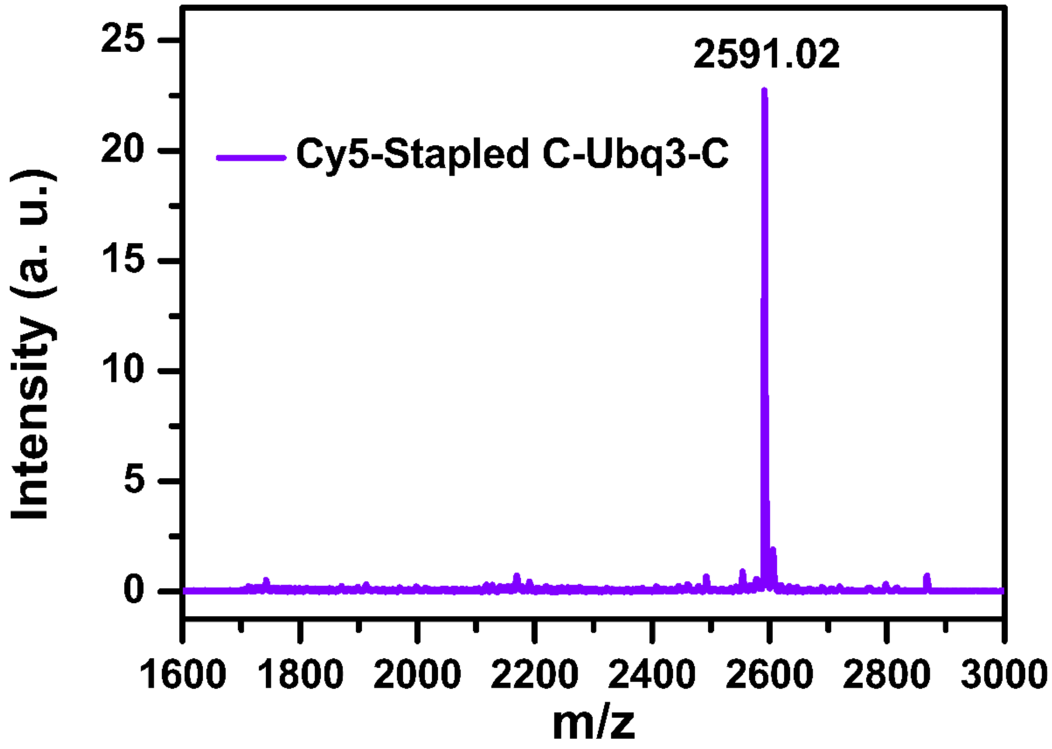


**Figure S8.** MALDI Mass Spectrum of the Cyanine5-labelled Stapled Cys-Ubq3-Cys Peptide (Peptide sequence: **Cy5-Cys-MQIFVKTLTGKTITLEV-Cys**).

9.

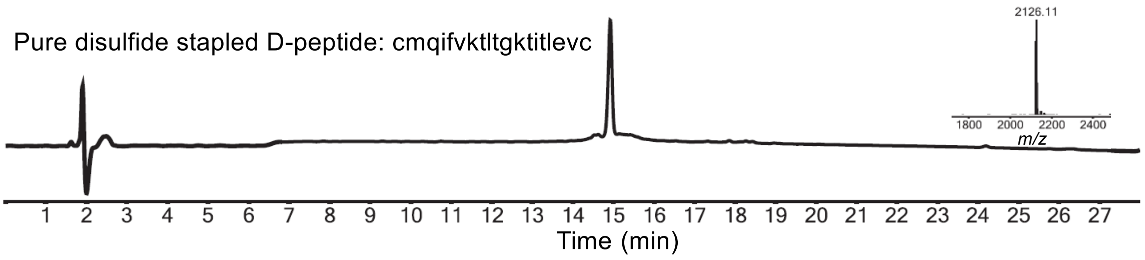


**Figure S9.** Analytical RP-HPLC profile ($\lambda$ = 214 nm) together with ESI-MS data (inset) of purified oxidized D-Ubiquitin peptide segment. Linear gradient, 20-64% of B over 22 min including 4 min equilibration time using Agilent Zorbax SB-C3, 5 µm 4.6 x 150 mm, LC column with 0.9 mL/min flow rate, was used for the chromatographic separation. Purification was performed using a linear gradient 25%-50% buffer B in buffer A over 50 min with a flow rate of 5 mL/min at 40 °C (buffer A = 0.1% TFA in water; buffer B = 0.08% TFA in acetonitrile) using a C4, 10 x 250 mm, preparative HPLC column (Phenomenex proteo, 300 Å, 10 µm).

10.

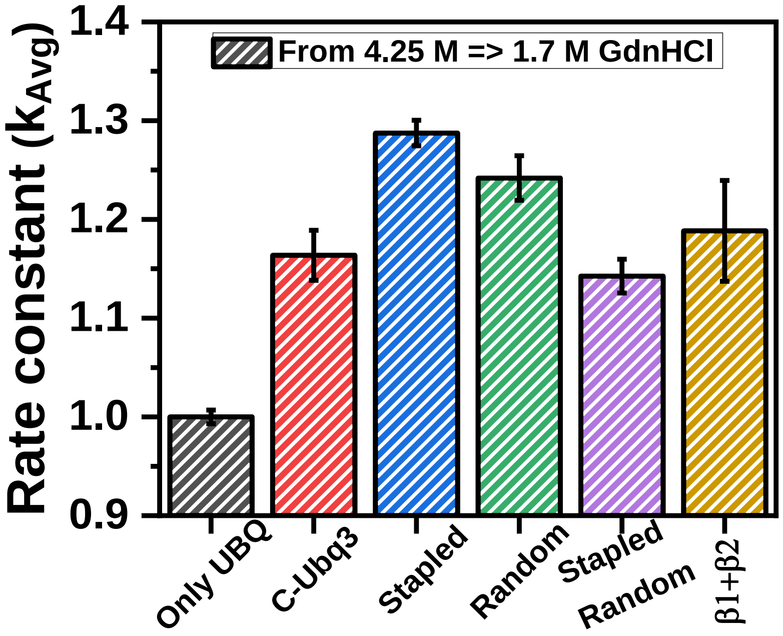


**Figure S10.** Normalized folding rate constants of Ubiquitin alone, in presence of the stapled decoy peptide and all the control peptides at a final GdnHCl concentration 1.7 M GdnHCl from a starting GdnHCl concentration of 4.25 M, protein in 4.25 M GdnHCl (30 µM stock concentration) and peptides in buffer (200 µM stock concentration, except β1 + β2).

11.

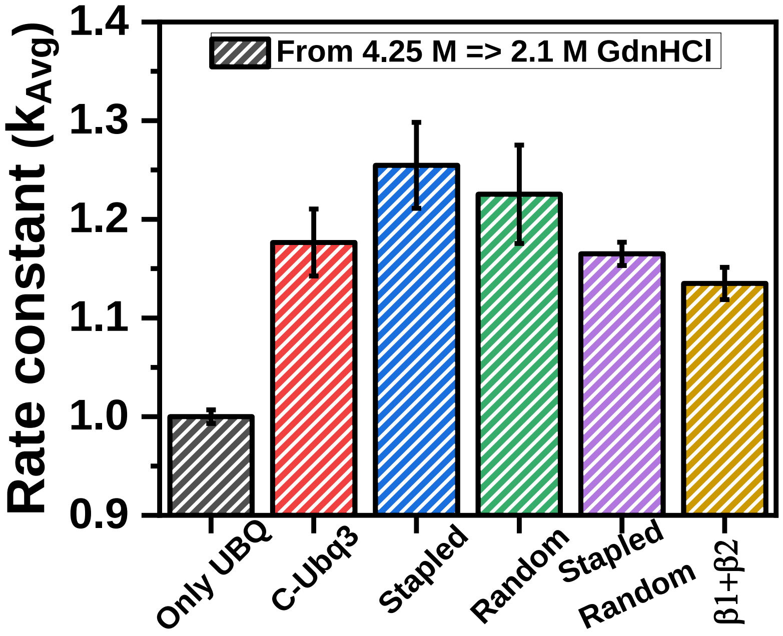


**Figure S11.** Normalized folding rate constants of Ubiquitin alone, in presence of the stapled decoy peptide and all the control peptides at a final GdnHCl concentration 2.1 M GdnHCl from a starting GdnHCl concentration of 4.25 M, protein in 4.25 M GdnHCl (30 µM stock concentration) and peptides in buffer (200 µM stock concentration, except β1 + β2).

12.

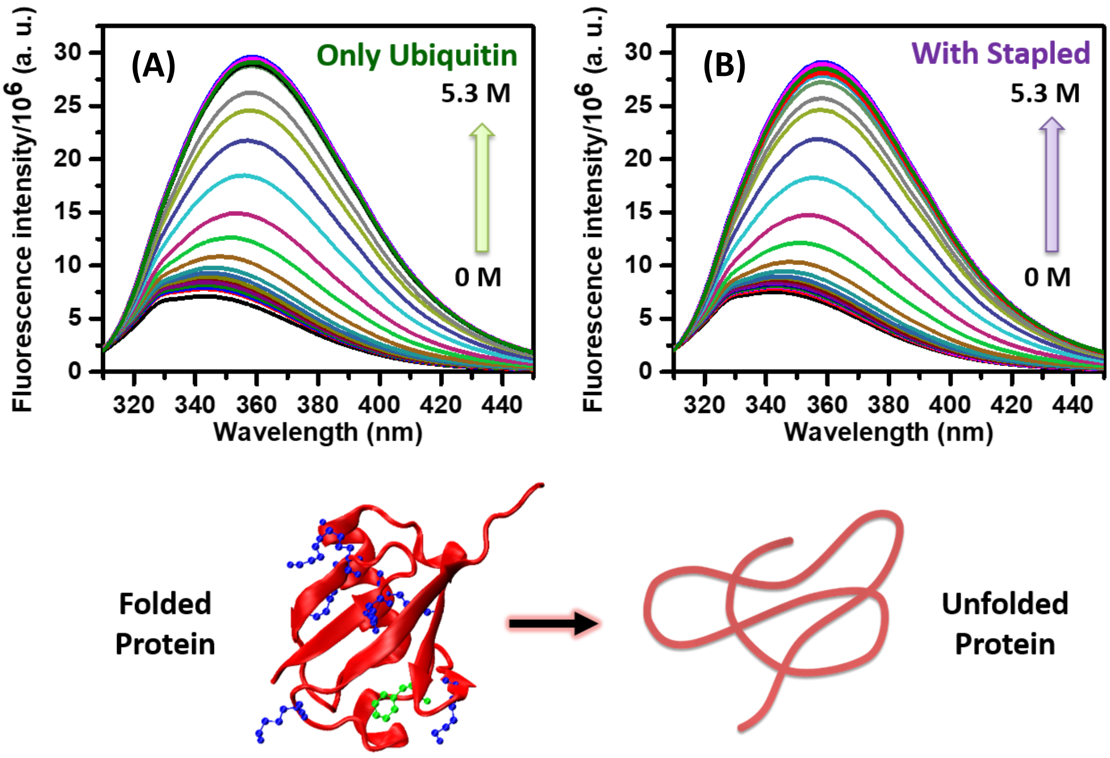


**Figure S12.** Titration (Steady-state Fluorescence Spectra) of Ubiquitin F45W (10 µM) with increasing concentration of GdnHCl (A) in absence and (B) in presence of the stapled decoy peptide (40 µM), Cys-Ubq3-Cys measured by monitoring fluorescence at 360 nm.

13.

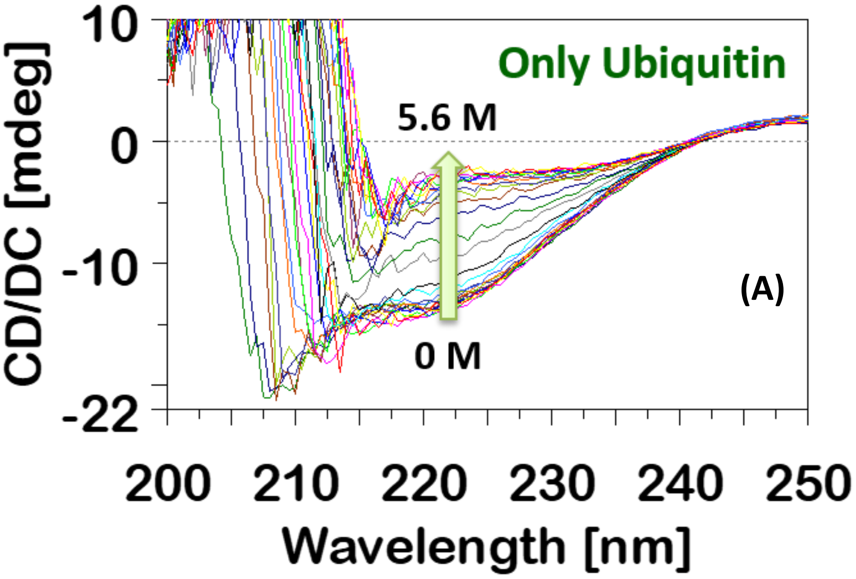

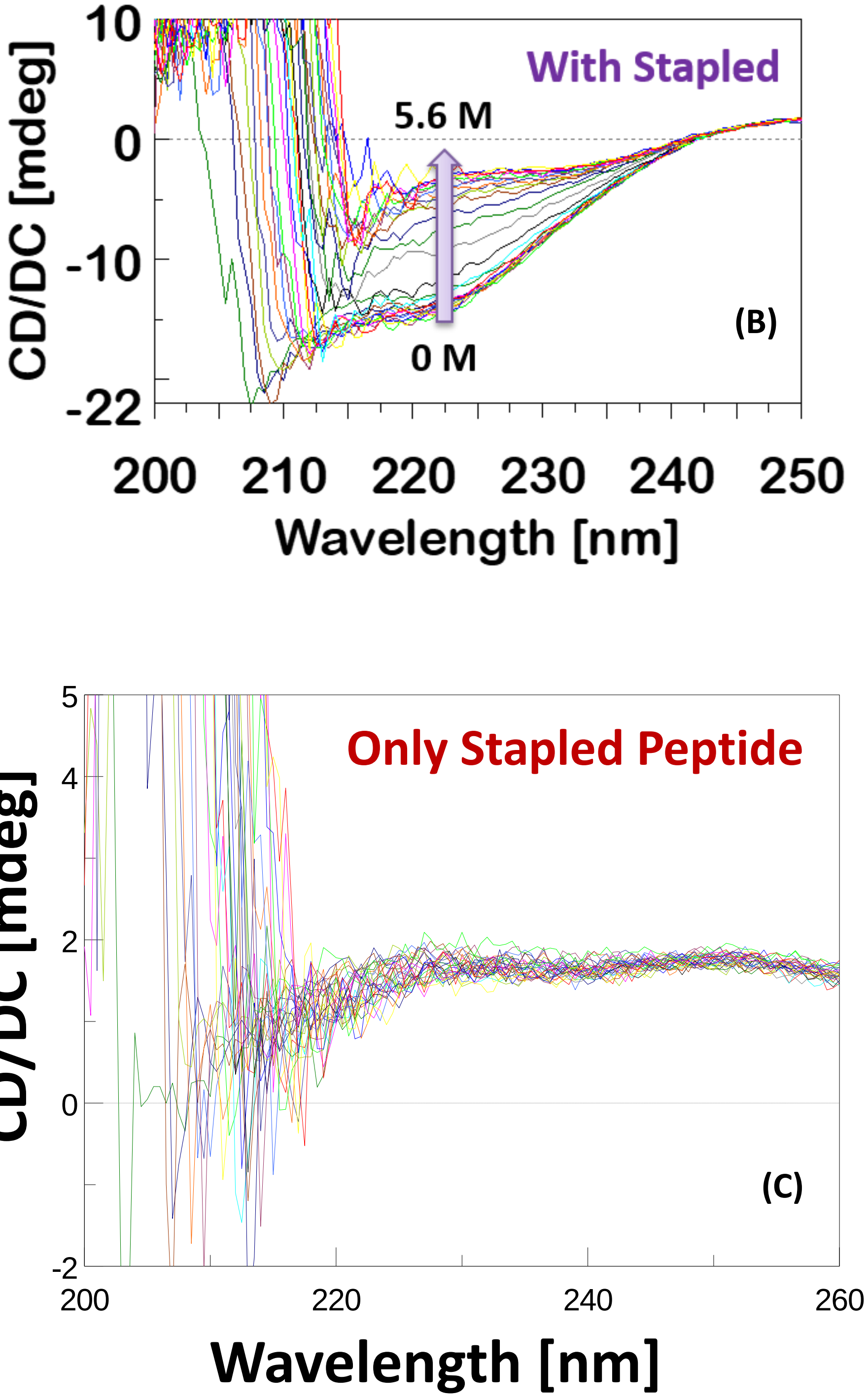


**Figure S13.** Titration (Steady-state denaturation) of Ubiquitin F45W (10 μM) with increasing concentration of GdnHCl (A) in absence and (B) in presence of the stapled xenonucleus peptide (40 μM), Cys-Ubq3-Cys measured by monitoring circular dichroism (CD) from λ = 200 - 250 nm. (C) CD of stapled Cys-Ubq3-Cys (40 μM) alone at those above different GdnHCl concentrations.

14.

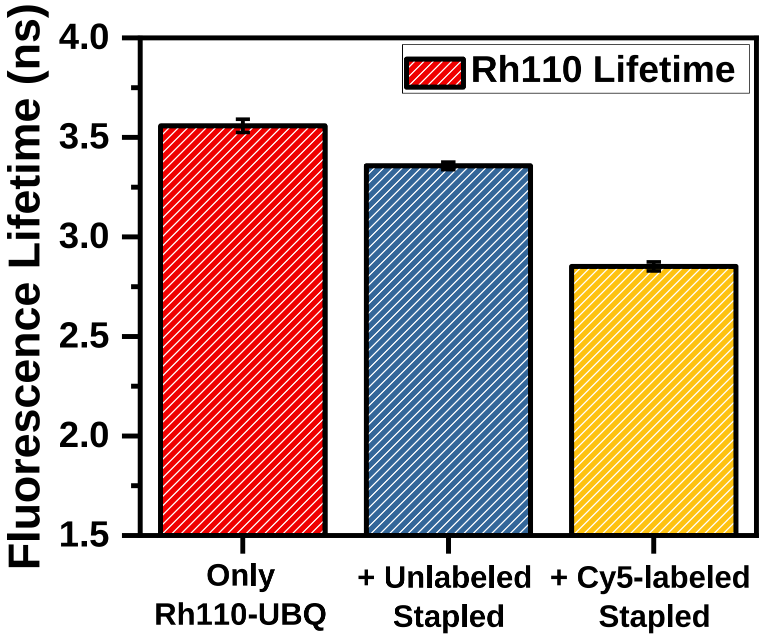


**Figure S14.** Fluorescence lifetimes ($\tau_{Avg}$) of Rhodamine 110 in only 100 nM Rh110-labelled Ubiquitin (Red), and in presence of (~120 µM) unlabelled Stapled peptide (blue), and in presence of (~3 µM) Cy5-labelled Stapled peptide (yellow); all in 3 M final GdnHCl concentration. Here, $\tau_{Avg}$ is the intensity averaged lifetime, i.e. $\tau_{Avg} = \frac{\sum_{i=1}^{2}\alpha_i \tau_i^2}{\sum_{i=1}^{2}\alpha_i \tau_i}$ , '$\alpha_i$' is the amplitude and '$\tau_i$' is the corresponding lifetime (in ns) of the i-th component.

15.

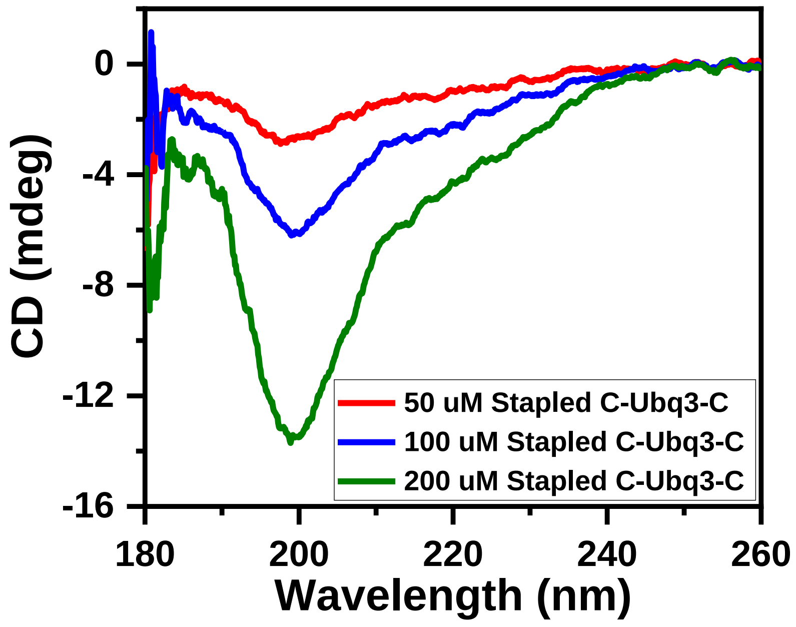

**Figure S15.** Circular dichroism spectra (far-UV CD) of the xenonucleus (stapled Cys-Ubq3-Cys peptide) in water, at the concentrations of 50 μM (Red), 100 μM (blue) and 200 μM (green), respectively.

16.

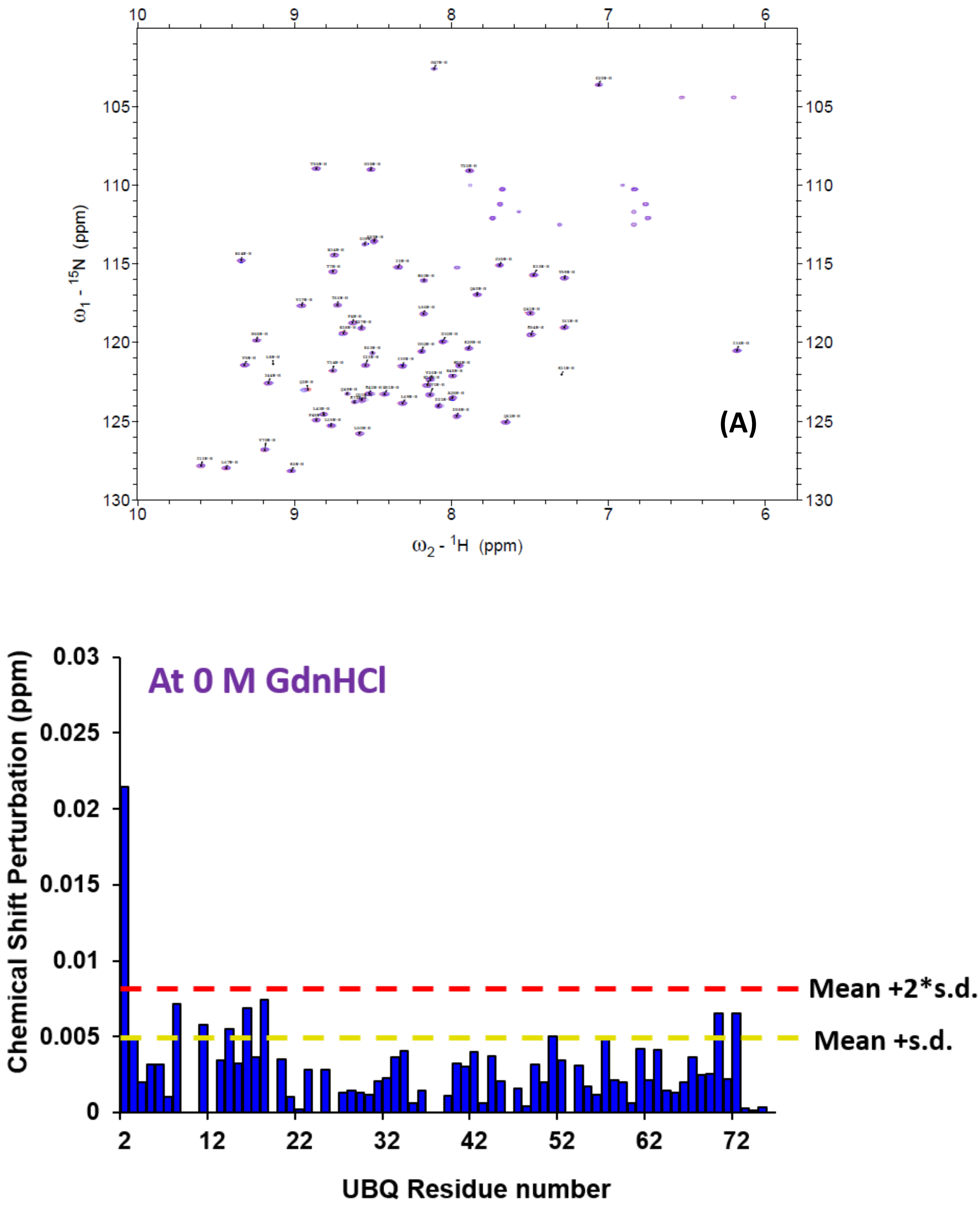


**Figure S16.** (A) Overlay of the HSQCs of free UBQ (blue) and UBQ/Stapled Xenonucleus complex (red) at 0 M GdnHCl. (B) Chemical Shift Perturbation [((dN/5)^2+(dH)^2))$^{1/2}$] between UBQ and UBQ/peptide (Stapled L-peptide) complex at 0 M GdnHCl.

17.

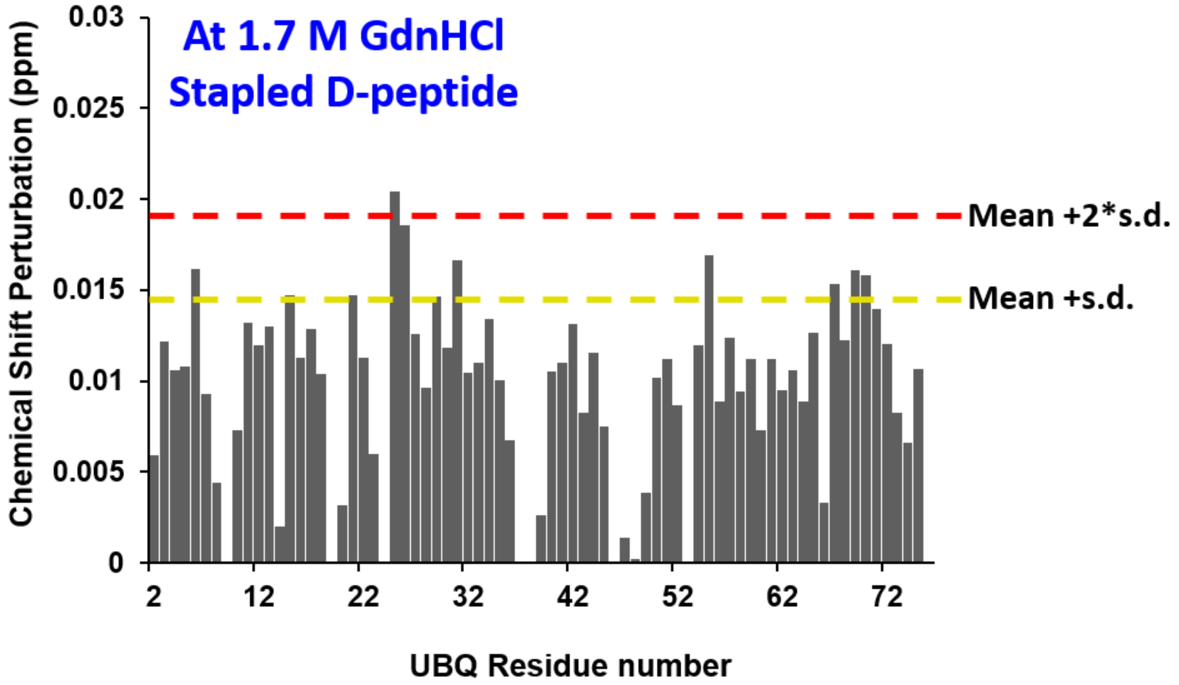


**Figure S17.** Chemical Shift Perturbation $[((dN/5)^2+(dH)^2))^{1/2}]$ between Ubiquitin (UBQ) and UBQ/peptide (Stapled D-peptide) complex at 1.7 M GdnHCl.

18.

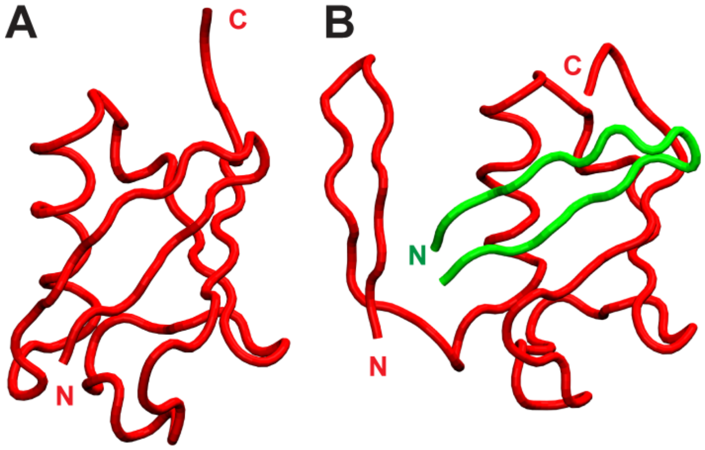


**Figure S18.** Representative structures from the M and M' minima. N and C-termini are marked. These figures were drawn using tube representation in VMD[1] (A) Structure of the folded UBQ monomer from the minimum marked as M in the Fig. 5E (B) Structure of the UBQ in the M' state, which is seen as an extra dip in the free energy plot apart from U and M. Only the simulations with stapled peptide accumulate considerable amount of the M' state. This state corresponds to the protein UBQ folded on the extra β1-β2 (colored in green) instead of its own β1-β2, which itself is folded but is just floating around.

19.

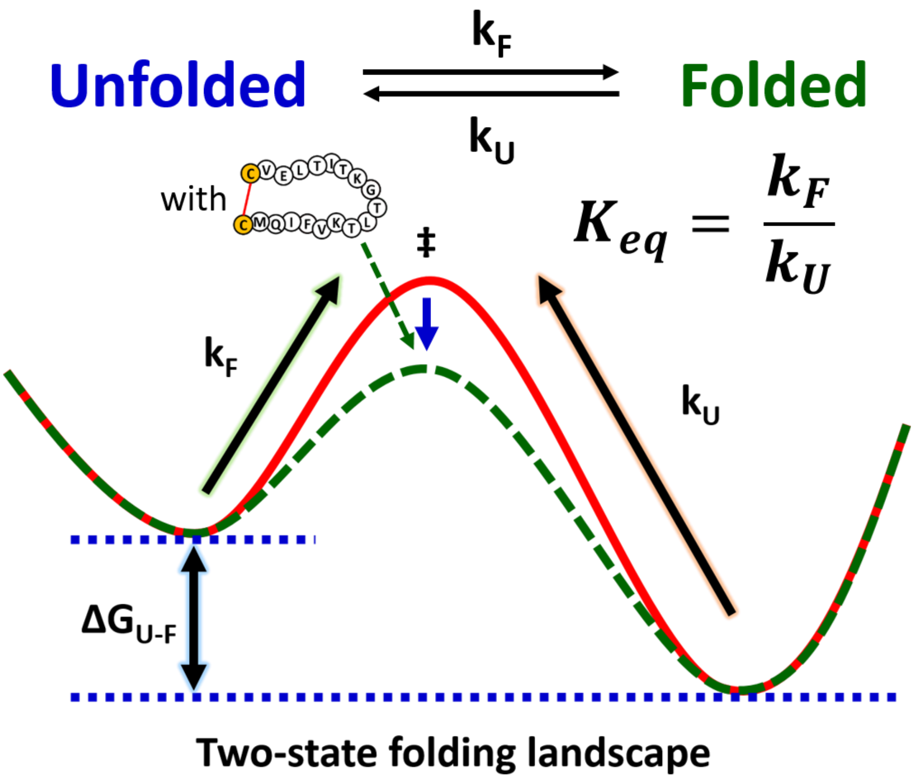


**Figure S19.** Hypothetical thermodynamic model for the protein-xenonucleus interaction. The xenonucleus works like a ‘mini folding-catalyst’.

| | No. of transitions UBQ + Unstapled Xenonucleus | No. of transitions UBQ + Stapled Xenonucleus |
|---|---|---|
| M’ => M | 138 ( 22%) | 440 ( 14%) |
| M’ => U | 478 ( 78%) | 2690 ( 86%) |
| Total transitions | 616 | 3130 |

**Table S1.** The population distribution of the trajectories for simulations of UBQ with unstapled and stapled xenonucleus. Trajectories initiating from the M' state and ending in either U or M state were calculated.

| Steady-state Fluorescence Experiment | | Value | Standard Error |
|---|---|---|---|
| Fitting Equation | y = (A+B*exp(-m*0.00168*(c-x)))/(1+exp(-m*0.00168*(c-x))) | | |
| Only Ubiquitin | A | 6.93E6 | 140375.93 |
| | B | 2.92E7 | 167794.25 |
| | m (cal/mol-M) | 1922.05 | 80.25 |
| | c (in M) | 3.00 | 0.01 |
| | $\Delta G^0_{unf}$ (kcal/mol) | 5.77 | 0.24 |
| Ubiquitin with xenonucleus | A | 6.87E6 | 126910.37 |
| | B | 2.83E7 | 149353.38 |
| | m (cal/mol-M) | 2029.08 | 81.35 |
| | c (M) | 2.98 | 0.01 |
| | $\Delta G^0_{unf}$ (kcal/mol) | 6.05 | 0.24 |

**Table S2.** Parameters extracted from the equilibrium chemical denaturation experiments (using steady-state fluorescence) for only ubiquitin (F45W) and ubiquitin with the stapled xenonucleus. The parameters were obtained by fitting the data given in Figure 2 (A) with the sigmoid function mentioned in the manuscript.

| Circular Dichroism Experiment | | Value | Standard Error |
|---|---|---|---|
| Fitting Equation | y = (A+B*exp(-m*0.00168*(c-x)))/(1+exp(-m*0.00168*(c-x))) | | |
| Only Ubiquitin | A | -13.30 | 0.12 |
| | B | -3.37 | 0.14 |
| | m (cal/mol-M) | 2116.62 | 178.26 |
| | c (M) | 3.16 | 0.03 |
| | $\Delta G^0_{unf}$ (kcal/mol) | 6.69 | 0.57 |
| Ubiquitin with xenonucleus | A | -14.06 | 0.17 |
| | B | -3.71 | 0.20 |
| | m (cal/mol-M) | 2094.51 | 240.38 |
| | c (M) | 3.12 | 0.04 |
| | $\Delta G^0_{unf}$ (kcal/mol) | 6.53 | 0.75 |

**Table S3.** Parameters extracted from the equilibrium chemical denaturation experiments (using steady-state circular dichroism) for only ubiquitin (F45W) and ubiquitin with the stapled xenonucleus. The parameters were obtained by fitting the data given in the inset of Figure 2 (A) with the sigmoid function mentioned in the manuscript.

| Residue | Nearest Neighbour(s) {through space} |
|---|---|
| Val26 | 1. Val17 (4.24 Å between the terminal carbon of both valines)<br>2. Ile3 (4.92 Å between the terminal carbons) |
| Lys29 | Glu16 (2.67 Å between terminal Nitrogen of Lys and backbone oxygen of Glu) |
| Lys33 | Thr14 (3.407 Å between terminal nitrogen of lysine and backbone oxygen of threonine) |
| Leu67 | 1. Val5 (4.651 Å between terminal carbons of both)<br>2. Ile3 (3.871 Å between terminal carbons of both) |
| Leu69 | Thr7 (4.20 Å between terminal carbon of both) |
| Leu71 | 1. Thr9 (7.22 Å between terminal carbons)<br>2. Leu8 (6.87 Å between terminal carbon of 8 and backbone carbon of 71) |

**Table S4.** Nearest neighbours (through space) and their distances for residues with maximum CSPs, based on PDB ID 1UBQ. Lys29-Glu16 (helix-beta2 interaction) is the closest followed by Lys33-Thr14. Both have a potential to form salt bridge interactions followed by the hydrophobic interactions between val26-val17, Leu67-Ile3, Leu69-Thr7, respectively.

| Sample | $\alpha_1$ | $T_1$ | $\alpha_2$ | $\tau_2$ | $\tau_{Avg}$ |
|---|---|---|---|---|---|
| Only Rh110-UBQ | 9421.87 ± 940.78 | 3.65 ± 0.05 | 1398.9 ± 277.85 | 0.86 ± 0.17 | 3.56 ± 0.03 |
| Rh110-UBQ + Unlabelled Xenonucleus | 11055.2 ± 1445.14 | 3.51 ± 0.03 | 2512.12 ± 493.36 | 1.03 ± 0.19 | 3.36 ± 0.02 |
| Rh110 + Cy5-labelled Xenonucleus | 10680.53 ± 3662.80 | 3.10 ± 0.03 | 6701.67 ± 2058.25 | 0.52 ± 0.02 | 2.85 ± 0.02 |

**Table S5.** Analysis of the fluorescence lifetime traces of Rh110-labelled ubiquitin alone, and when incubated with the unlabelled xenonucleus and the Cy5-labelled xenonucleus.
